# The role of recent speciation in present-day patterns of tetrapod phylogenetic relatedness

**DOI:** 10.1101/2023.11.03.565445

**Authors:** Hector Tejero-Cicuendez, Iris Menendez, Elizabeth M. Steell, Guillermo Navalon, Fernando Blanco, Jiri Smid

## Abstract

**Aim:** Biodiversity is distributed unevenly among lineages and regions, and understanding the processes generating these global patterns is a central goal in evolutionary research, particularly in light of the current biodiversity crisis. Here, we integrate phylogenetic relatedness with species diversity patterns in four major clades of living tetrapods (amphibians, squamates, birds, and mammals) to approach this challenge.

**Location:** Global.

**Time period:** 300 million years ago – Present.

**Major taxa studied:** Tetrapods.

**Methods:** We studied geographic patterns of richness-corrected phylogenetic diversity (residual PD), identifying regions where species are phylogenetically more closely or distantly related than expected by richness. We explored the effect of different factors in residual PD: recent speciation rates, temporal trends of lineage accumulation, and environmental variables. Specifically, we searched for evolutionary and ecological differences between regions of high and low residual PD.

**Results:** Our results show heterogeneous spatial patterns of diversity dynamics across tetrapods. They reveal an overall negative relationship between recent speciation rates and residual PD, underscoring the role of recent speciation events in structuring current biogeographic patterns. Furthermore, we found differences between endothermic and ectothermic tetrapods in response to temperature and precipitation, highlighting the pivotal role of thermal physiology in shaping diversity dynamics.

**Main conclusions:** Geographic patterns of diversity dynamics are heterogeneous across tetrapod clades and help us disentangle the evolutionary and ecological processes underlying them. By illuminating the multifaceted factors underpinning global diversity patterns, our study represents a significant advancement towards better understanding of how the present-day diversity of tetrapods was formed and how speciation rates influenced their species and phylogenetic diversity across clades and regions.

## Introduction

Clarifying the evolutionary and ecological processes underlying present-day patterns of biodiversity remains a central goal in evolutionary biology (de Candolle 1859; Matthew 1915; Moore 1920; Ruthven 1920; Allee 1926; Dobzhansky 1950; Fischer 1960; MacArthur 1965; Anderson 1974; Harmon 2012; Saupe 2023). Even though the study of global geographic patterns of species richness has occupied a prominent role in macroecological discussion since the infancy of evolutionary biology (Humboldt and Bonpland 1807; Darwin 1859; Wallace 1876), it is only in recent decades that the discipline has addressed the importance of accounting for the phylogenetic relationships among species (Nee and May 1997; Pyron and Wiens 2013; Wiens 2015). The emergence and development of spatial phylogenetics (Earl et al. 2021; Mishler 2023) has advanced our understanding of macroecological dynamics by combining phylogenetic relatedness with geographic biodiversity patterns. Specifically, the use of phylogeny-based metrics of biodiversity, such as Faith’s phylogenetic diversity (PD; Faith 1992), enables investigations into the geographic distribution of species relatedness by considering the length of the phylogenetic branches connecting all the species present in a region. For instance, high PD values indicate the sympatric presence of distantly related species, while low PD values result from closely related species inhabiting a given geographic area. The explicit inclusion of the phylogenetic dimension may greatly improve our ability to elucidate the synergistic effects of evolutionary, ecological, and environmental factors on diversity dynamics (Davies and Buckley 2011; Tucker et al. 2017) in addition to providing an essential source of information for conservation purposes (Faith 1992; Redding and Mooers 2006).

In most cases, species richness and PD are positively correlated: regions showing high and low species richness have high and low levels of PD, respectively (e.g., Davies and Buckley 2011; Fritz and Rahbek 2012). A positive linear relationship between richness and PD is expected under a null scenario of balanced phylogeny and species distributions: if all the species and clades present in particular regions were subjected to homogeneous and constant diversification and dispersal rates, a change in the number of species would be reflected in a proportional change in the number of clades, such that the degree of phylogenetic relatedness would vary accordingly and invariably across regions. This makes species richness a generally good proxy of PD (Rodrigues et al. 2005; Morlon et al. 2011). However, geographic patterns of richness and PD are not necessarily congruent (Tucker and Cadotte 2013). In other words, there are geographic regions where species are more distantly (high PD) or more closely (low PD) related than would be predicted by the number of species that these regions harbor (Fig. 1a). These deviations from the expected relationship of PD to species richness (residual PD; Velasco and Pinto-Ledezma 2022) are the result of variations in the generative processes of biodiversity: speciation, extinction, and dispersal (Ricklefs 2004; Mittelbach et al. 2007; Wiens 2011). Therefore, studying such variations across regions and clades is essential to understand the evolutionary role of these processes, as well as the biotic and abiotic factors underpinning geographic patterns of biodiversity.

**Fig. 1.**
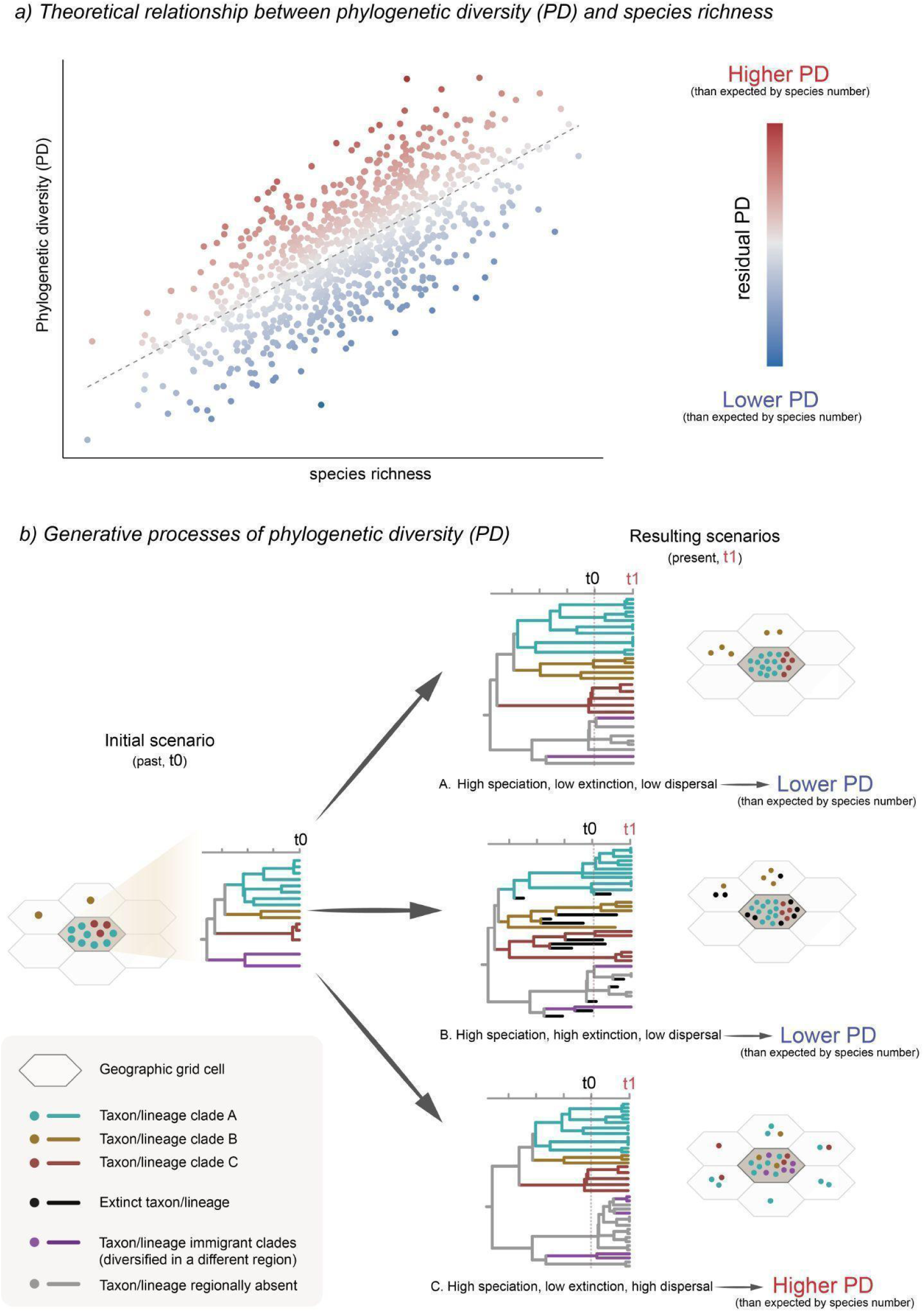
Theoretical conceptualization of this study. of a) Schematic representation of the relationship between species richness (X-axis) and phylogenetic diversity (Y-axis). Each point represents a geographic grid cell with color representing residual phylogenetic diversity: in red, the cells with more distantly related species (high PD) and in blue more closely related (low PD) than would be predicted by the number of species that these regions harbor. b) Hypothetical scenarios with different levels of PD produced by different combinations of speciation, extinction, and dispersal.

For example, high residual PD values might represent the so-called “museums” or “sanctuaries” of biodiversity (Dobzhansky 1950; Stebbins 1974) resulting from the gradual accumulation of species, either by dispersal from other areas (immigration) or by low levels of speciation and extinction (low turnover rate), but they might also arise owing to a different combination of processes, such as high speciation rates in the past followed by low extinction rates of old lineages, or, alternatively, exceptionally high extinction rates of younger lineages. Conversely, low residual PD can indicate “cradles” of biodiversity (Stebbins 1974) resulting from high speciation and extinction rates maintained through time (high turnover rate), but it can also arise in other scenarios, such as reduced extinction rates of young lineages or increased extinction of older clades. The multiplicity of scenarios able to generate similar patterns of lineage and phylogenetic diversity (e.g., Fig 1b) highlights the importance of investigating the underlying evolutionary processes beyond solely focusing on elucidating whether specific geographic regions are cradles or museums of biodiversity (Vasconcelos et al. 2022).

The increasing availability of both global distribution and phylogenetic data from species-rich clades allows for the exploration of large-scale diversity patterns and evolutionary processes. For example, in the last decade, the geographic distribution of residual PD has been addressed for the four major clades of living terrestrial vertebrates: mammals (Davies and Buckley 2011), amphibians (Fritz and Rahbek 2012), birds (Voskamp et al. 2017), and squamates (Gumbs et al. 2020; Vásquez-Restrepo et al. 2023). This wealth of data presents an exciting, but unrealized, opportunity for a detailed comparison of geographic patterns of residual PD across the four major clades of tetrapods, including critically identifying differences and similarities in the biotic and abiotic factors underpinning global patterns of tetrapod biodiversity. In this study, we characterize geographic patterns of tetrapod species and phylogenetic diversity, and we test hypotheses related to the impact of multiple factors on regional levels of residual PD: recent speciation rates, evolutionary time, and environmental conditions. Specifically, our goals are: 1) To identify similarities and differences in the geographic patterns of residual PD across tetrapod clades. This might allow for an informed subsequent investigation on the different processes underlying these patterns. 2) To test the role of recent speciation rates in shaping the observed geographic patterns. An intuitive hypothesis is that regions of low residual PD arise due to the sympatry of lineages with increased speciation rates, especially recent rates, but other factors (e.g., extinction, dispersal) also might impact geographic differences across clades. The fact that recent speciation rates can be reliably estimated from extant timetrees (Title et al. 2019; Louca and Pennell 2020) allows us to analyze the relative role of this process in generating the observed patterns. Importantly, regional residual PD and lineage speciation rates are calculated from different phylogenies (PD from regional phylogenies, speciation from complete phylogenies). 3) To explore differences in the temporal patterns of phylogenetic bifurcations and lineage accumulation between regions of high and low residual PD. We hypothesize that regional phylogenies of low residual PD areas will have a more recent accumulation of lineages and speciation events than those of high residual PD areas, which would concentrate lineage origination toward the root of the clade. 4) To test the relationship of present-day environmental variables with current geographic patterns of residual PD, focusing on potential environmental differences between regions of exceptionally high and low residual PD. Interestingly, this may reveal fundamental differences in the evolutionary dynamics of different tetrapod clades, and may further help test hypotheses about the interplay of evolutionary and historical factors in shaping their biodiversity patterns.

## Materials and Methods

### Vertebrate data

We obtained distribution vector data for amphibians and terrestrial mammals from IUCN (IUCN 2022), for birds from BirdLife International (http://www.birdlife.org/) and for squamates from (Roll et al. 2021). Range maps for all groups were downloaded on 1 May 2022. Phylogenetic data for all groups were downloaded from VertLife (https://data.vertlife.org/). This includes the consensus and posterior phylogenetic trees for amphibians (Jetz and Pyron 2018), birds (Jetz et al. 2012; backbone from Hackett et al. 2008), mammals (Upham et al. 2019), and squamates (Tonini et al. 2016). After matching both distribution and phylogenetic data, the final datasets contained a total of 28,270 species: 5832 amphibians, 7995 birds, 5164 mammals, and 9279 squamates. These were the datasets used in subsequent analyses.

### Geographic grid and species richness

For each group, we first produced a hexagonal 100-km-resolution species richness grid using the epm package v1.1.1 (Title et al. 2022) in R 4.3.0 (R Core Team 2023), with the polygon distribution data transformed into an equal-area Behrmann projection as input and the ‘centroid’ approach. The resulting grid contains the information of the species present in each hexagonal cell, and was the base cell grid for all subsequent analyses.

### Residual phylogenetic diversity

We obtained an average phylogenetic diversity (PD) grid after calculating PD grids for 100 trees from the posterior distribution for each tetrapod group, accounting for phylogenetic (temporal and topological) uncertainty. These grids were produced with the functions addPhylo and gridMetrics in epm (Title et al. 2022), and they represent the sum of the branch lengths of the phylogenetic tree connecting all species in each cell (Faith’s PD; Faith 1992). We used the R package rlist v0.4.6.2 (Ren 2021) to process the ‘posterior’ grids and ultimately obtain a grid of average PD values. With the per-cell values of species richness and PD, we performed a local regression analysis (LOESS) with the smoothing parameter α = 0.75 and obtained the residuals from it. We then mapped these residuals again onto the original hexagonal grid to visualize the geographic distribution of the deviation of PD relative to richness (residual PD). High residual values indicate high PD for a given number of species (i.e., the species within a grid cell are more distantly related to each other than expected by the species richness of the grid cell), and, conversely, low residual PD indicates that the species present in a grid cell are more closely related to each other than predicted by species richness. To identify focal regions of particularly extreme values of residual PD, we set a threshold at 10% (i.e., lowest residual PD) and 90% (i.e., highest residual PD) from the total distribution of values from each vertebrate clade. We then visually identified focal regions of interest with elevated density of contiguous high and low residual PD grid cells to investigate whether there are differences between them in speciation rates, lineage accumulation patterns, and environmental conditions.

### Recent speciation rates

We estimated recent speciation rates (tip rates) calculating the average DR metric (Jetz et al. 2012) across 100 trees from the posterior distribution for each vertebrate clade, accounting for phylogenetic uncertainty in these speciation estimates. Then, we calculated mean DR values for each hexagonal grid cell (mean DR values of the species present in each grid cell). We conducted a linear regression model of per-cell mean DR against residual PD (see above). To further understand the links between speciation rates and patterns of geographic diversity, we additionally tested for differences in DR values between species present in regions of highest and lowest residual PD. To do this, we performed linear models with randomized residual permutations with the RRPP package v1.3.1 (Collyer and Adams 2018, 2022), first to globally compare grid cells of high and low residual PD and then to individually compare among the focal regions we identified (see above).

### Patterns of lineage accumulation and speciation events

We searched for differences in lineage accumulation patterns between regions of high and low residual PD by calculating the number of lineages through time for extreme regions falling within the 10% lowest and highest (i.e., 10% and 90% thresholds) values of residual PD using the R package ape v5.7.1 (Paradis and Schliep 2019), with the aid of geiger v2.0.11 (Pennell et al. 2014) and phytools v1.5.1 (Revell 2012) for phylogenetic data handling. Visual inspection of lineage-through-time (LTT) plots allows us to determine whether our data reflect two main expectations: i) regions of high residual PD should exhibit comparatively older lineages than regions with low residual PD values, which might indicate that these regions acted as reservoirs of ancestral diversity, and ii) the pattern of lineage accumulation might be different between regions of high and low residual PD. For each of these regions of extreme (high and low) levels of residual PD, we calculated the gamma statistic (Pybus and Harvey 2000) on each of the 100 posterior trees pruned to represent regional phylogenies. The gamma statistic informs of deviations in the concentration of speciation events from a pure birth model toward the present (gamma > 0; internal nodes closer to the tips) or toward the past (gamma < 0; internal nodes closer to the root). This allows us to test whether differences between extremely high and low residual PD are related to temporal differences in the occurrence of speciation events. The gamma-statistic was implemented through the function *gammaStat* in the package ape v5.7.1 (Paradis and Schliep 2019).

### Environmental variables

One of the factors that may affect evolutionary processes and therefore shape geographic patterns of biodiversity is the environment in which species live. We tested the relationship between residual PD and different environmental variables: mean annual temperature, temperature seasonality, annual precipitation, precipitation seasonality, net primary productivity (NPP), and terrain roughness index (TRI, a variable representing topographic complexity). Temperature and precipitation data were collected at a 10-minute spatial resolution (∼18.5 km) from the summary data for the period between the years 1970 and 2000 contained in the WorldClim v2.1 dataset (Fick and Hijmans 2017). Net primary productivity (NPP) data summarized over the period between 1981 and 2015 was obtained at 5-arc-minute resolution from the NDVI3g time series (Pinzon and Tucker 2014). The current topography data (Terrain Roughness Index, TRI) were based on (Wilson et al. 2007) and obtained from the ENVIREM dataset (Title and Bemmels 2018) at a spatial resolution of 10 arc-minutes. All the environmental variables were resampled to match the spatial resolution of the hexagonal cell grid built for species richness and phylogenetic diversity (100 km), so that we could have per-cell values for every variable in order to implement regression models. We independently tested the relationship between residual PD and the six environmental variables (precipitation, temperature, precipitation seasonality, temperature seasonality, NPP, and TRI), and we also tested the effect of the interaction of temperature and precipitation with a multiple regression model. Additionally, we generated three climatic spaces: one defined by temperature and precipitation, another one defined by temperature seasonality and precipitation seasonality, and a third one defined by NPP and TRI, and mapped the grid cells with lowest and highest 10% of the residual PD onto those climatic spaces to explore potential segregation between them. Finally, we also explored how residual PD is distributed across the latitudinal gradient, to compare residual PD patterns with species richness. Statistical analyses were performed with the RRPP package (Collyer and Adams 2018, 2022).

## Results

### Geographic patterns of residual phylogenetic diversity

We found some regions with consistent patterns of residual phylogenetic diversity (PD) across all four tetrapod clades (Supp. Fig. 1). The African continent harbors overall high residual PD (i.e., individual species more distantly related to each other than expected for the species richness of the assembly) for all the four clades, except for the Sahara Desert for mammals and the rainforest in central Africa for squamates. Conversely, large areas of South America contain low residual PD (species more closely related to each other than predicted by the species richness of the assembly) for all clades. In fact, Africa and South America constitute focal regions of highest and lowest residual PD, respectively, for all vertebrate clades (Fig. 2).

**Fig. 2.**
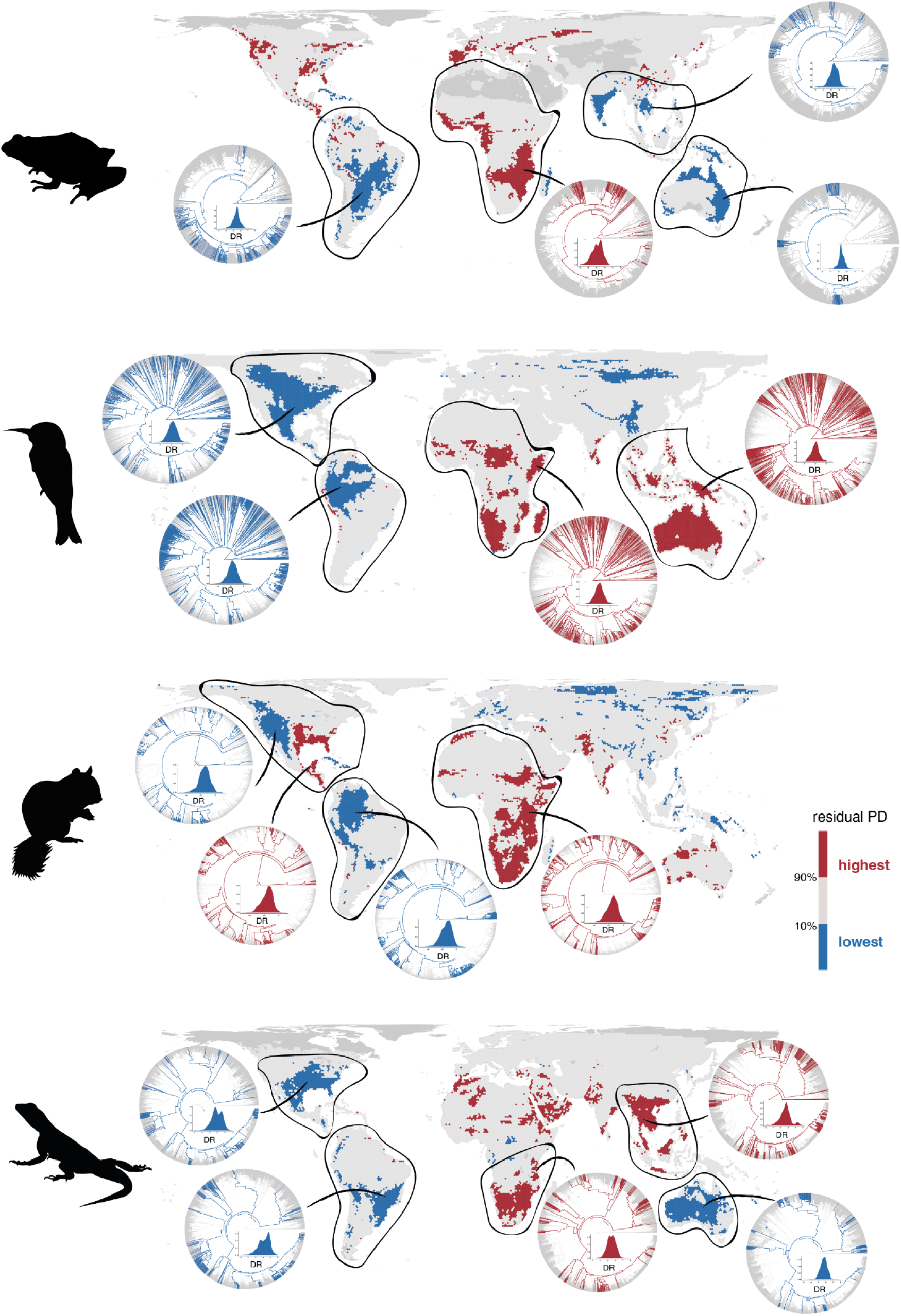
Geographic distribution of areas with the 10% lowest (in blue) and highest (in red) residual PD for terrestrial vertebrates (amphibians, birds, mammals, and squamates). The phylogenetic relationships of species present in focal regions, together with the density plot of recent speciation rates (DR) of those species, are also shown to illustrate the evolutionary differences of regions with high and low residual PD. Silhouettes by Guillermo Navalón and Sergio M. Nebreda.

On the other hand, some geographic regions exhibit very contrasting patterns of residual PD among the four clades. Australia is a low-residual-PD region for amphibians and especially squamates, while it harbors high residual PD for birds and mammals. Residual PD in the Indomalayan region is generally high for birds and squamates but relatively low for amphibians, while for mammals it is a very heterogeneous area with high levels in India but regions of low and high levels in Southeast Asia. North America is a region of low residual PD for birds and squamates, and of relatively high residual PD for amphibians. For mammals, there is a clear segregation in residual PD patterns between eastern (high values) and western (low values) North America. Eurasia is also a heterogeneous region across vertebrates, with generally low levels of residual PD for mammals and birds, and high levels for amphibians. The Arabian Peninsula contains exclusively high levels of residual PD for squamates, but for mammals and birds it shows relatively low levels across the interior and high levels in the mountainous regions of the south and west (Supp. Fig. 1).

### The effect of speciation rates

We found a statistically significant negative relationship between recent speciation rates (DR rates) and residual PD for all vertebrate clades (*P* <<< 0.001; Fig. 3). In other words, lineages exhibit, on average, higher recent speciation rates in regions of low residual PD (i.e., regions where species are more related to each other than predicted by the richness of the assembly), while regions of high residual PD generally have lower speciation rates. However, despite the strong significance, this negative relationship was found to be weak—due to the high dispersion of the data reflected in low R^2^ values—and variable among clades (Supp. Table 1; R^2^_amphibians_ = 0.007, R^2^_birds_ = 0.068, R^2^_mammals_ = 0.085, R^2^_squamates_ = 0.074).

**Fig. 3.**
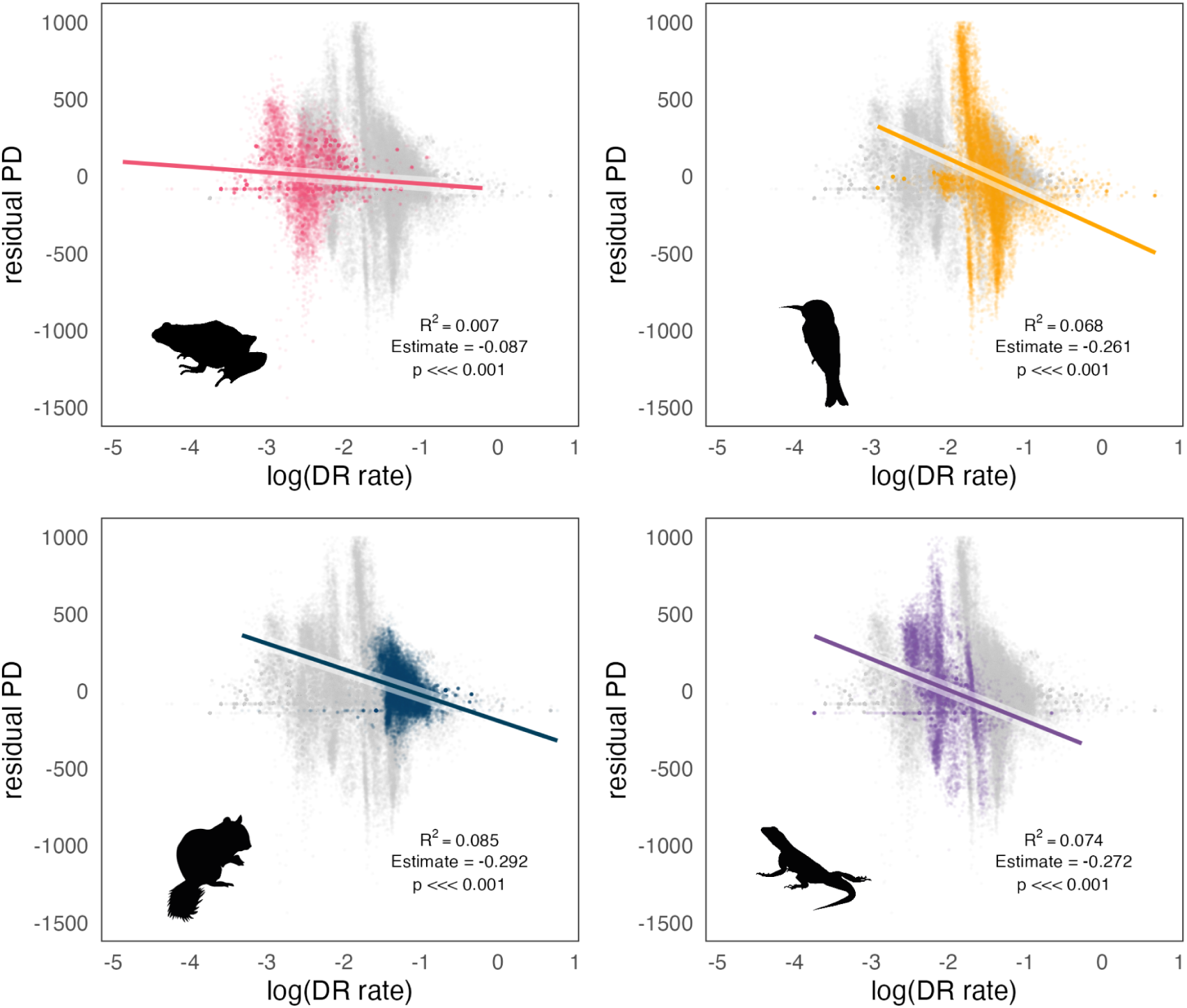
Relationship between recent speciation rates (DR rates) and residual PD for all four clades of terrestrial vertebrates. The grey cloud of points in the background of each plot shows results for the four vertebrate groups combined.

When comparing only regions falling in the 10% lowest and 10% highest residual PD, we found significantly lower speciation rates in the lineages present in grid cells with the lowest residual PD relative to those present in cells with the highest residual PD (*P* = 0.001 in all cases), especially for squamates, although, attending to the effect sizes (*Z*) and the overall distribution of DR values, this difference appears to be mild (Supp. Fig. 2).

Likewise, we found significant differences in species’ DR rates among individual focal regions of highest and lowest residual PD, with low residual PD broadly corresponding to higher speciation rates. Nonetheless, overall, these differences are not prominently apparent (Supp. Fig. 3, Supp. Table 2). Furthermore, we did not find clearly greater differences in DR effect size (*Z*) between focal regions of highest and lowest residual PD than between different regions of high residual PD and between different regions of low residual PD (*P* = 0.467; Supp. Fig. 4). This indicates that the differences in DR rates between regions of high and low residual PD, though significant, are not of particularly large magnitude.

### The effect of temporal patterns of lineage accumulation and speciation events

Our estimations of the gamma-statistic on the regional phylogenies show heterogeneous results (Fig. 4). In birds and squamates, regional species assemblages in the regions with lowest residual PD have overall gamma values above zero and higher than those in high residual PD regions, indicating a higher concentration of divergence events toward the present (i.e., in line with our working hypothesis that the sympatry of younger lineages results in low residual PD values). However, this difference between high and low residual PD regions is not so clear in mammals, for which assemblages in both kinds of regions show gamma values above zero. Furthermore, the pattern is the opposite in amphibians: low residual PD regions are related to gamma values below zero, and high residual PD regions show gamma values above zero (Fig. 4). This heterogeneity is evident in the per-region disaggregated results (Supp. Fig. 5), where there is no apparent relation between gamma values and the fact that regions have high or low residual PD.

**Fig. 4.**
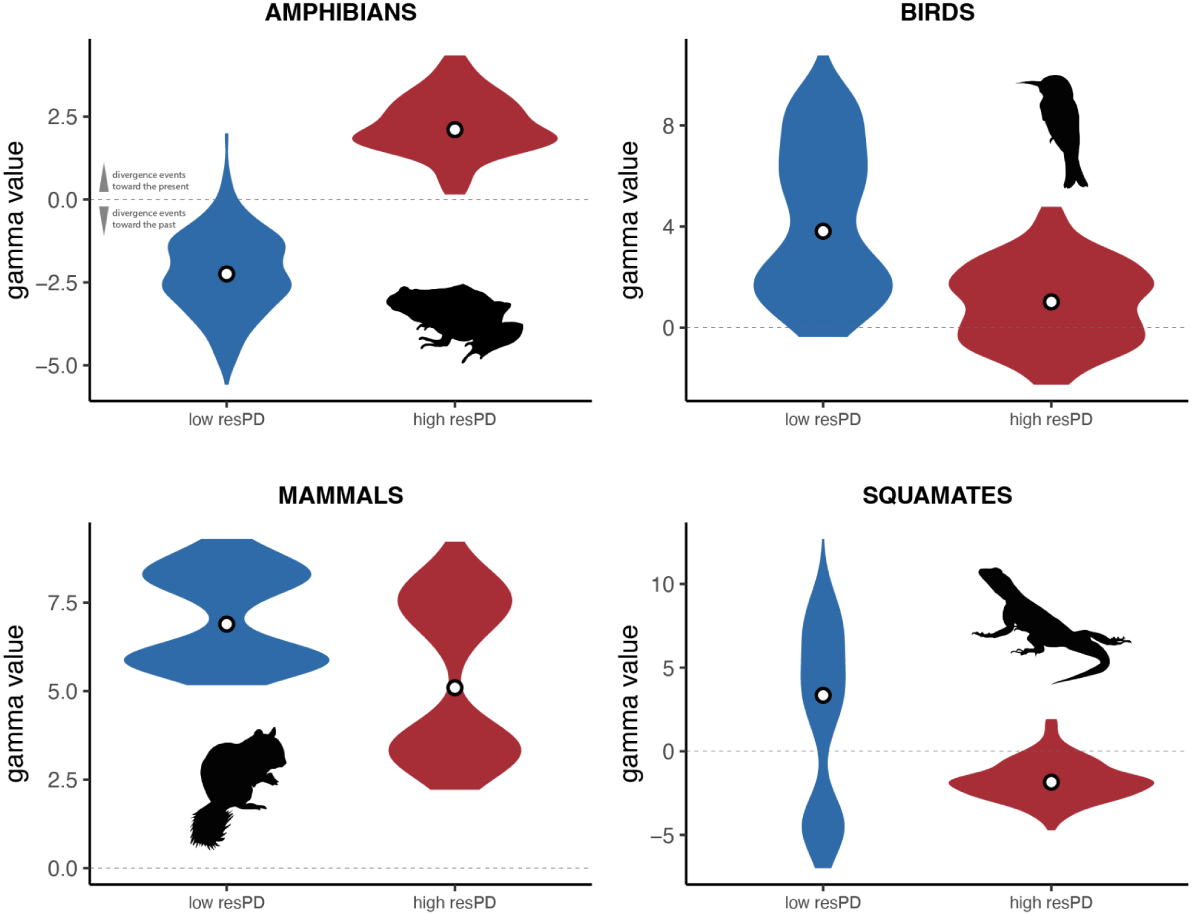
Gamma-statistic values in regions of lowest and highest residual PD. The gamma-statistic describes the temporal patterns of bifurcation events in the phylogeny: values above zero represent a bias toward more recent nodes than expected under a constant birth-death process, while gamma values below zero indicate that nodes are more concentrated to the past.

Similarly, we did not find notable differences between focal regions of high and low residual PD in the time of origin or the pattern of accumulation of the lineages they harbor (Supp. Figs. 6-9). Specifically, we did not find the lineages in regions of high residual PD to be as old as expected relative to those in low residual PD regions, and the pace of lineage accumulation is also not clearly distinguishable between regions of low and high residual PD. Even though some regions of low residual PD do harbor younger diversity than regions of high residual PD (e.g., amphibians in Oceania relative to amphibians in Africa, Supp. Fig. 6), this is the opposite in other cases (e.g., mammals in high residual PD Africa have younger ancestors than in low residual PD regions of North and South America, Supp. Fig. 8). In most cases, the age of origin of the lineages leading to present-day species and the trajectories of the accumulation curves are similar in regions of high and low residual PD.

### Environmental variables

We found an overall low to no linear relationship between residual PD and environmental variables in simple regression models (Supp. Figs. 10-15). There is a very weak negative relationship between annual precipitation and residual PD for all groups (Supp. Fig. 10), while the relationship with mean annual temperature is negative for amphibians and positive for the rest of vertebrate clades (Supp. Fig. 11), with mammals showing the highest amount of residual PD variance explained in both cases (precipitation R^2^_mammals_ = 0.017; temperature R^2^_mammals_ = 0.138). However, the results from multiple regression models including temperature and precipitation indicate that the interaction between the two is significant and a considerably larger portion of the residual PD variance is explained by their combined effects, especially in birds and mammals (R^2^_amphibians_ = 0.042, R^2^_birds_ = 0.161, R^2^_mammals_ = 0.225, R^2^_squamates_ = 0.094; Appendix I), as reflected in the environmental space defined by temperature and precipitation (see below). For precipitation seasonality and temperature seasonality, amphibians show an opposite trend to that of the other clades, although the variance explained is very low overall (Supp. Figs. 12 and 13). In amphibians, the correlation of residual PD with precipitation seasonality is negative (Supp. Fig. 12), whereas it is positive with temperature seasonality (Supp. Fig. 13). For the rest of vertebrates, these correlations are positive and negative, respectively. There is an extremely low correlation of residual PD with net primary productivity (NPP) for all clades (Supp. Fig. 14). With current topographic complexity (terrain roughness index, TRI), the correlation is also very low overall, but there is a clearer negative trend in birds and mammals (Supp. Fig. 15).

In the environmental space defined by mean annual temperature and annual precipitation, we found substantial overlap between regions of low and high PD for amphibians and squamates, but a clear segregation for birds and mammals (Fig. 5). In both birds and mammals, extremely low residual PD regions are characterized by two combinations: low precipitation with low to moderate temperature (which roughly correspond to areas of tundra, cold deserts and temperate grasslands; Whittaker 1975), and high precipitation with high temperature (i.e., tropical rainforest and savannah). Most of the high residual PD regions for birds and mammals, on the other hand, are found in environments with both low to moderate precipitation and high temperature (i.e., subtropical desert and savannah). However, highest residual PD regions for birds, unlike in mammals, are also found in areas with high precipitation and temperature (i.e., tropical rainforest and savannah).

**Fig. 5.**
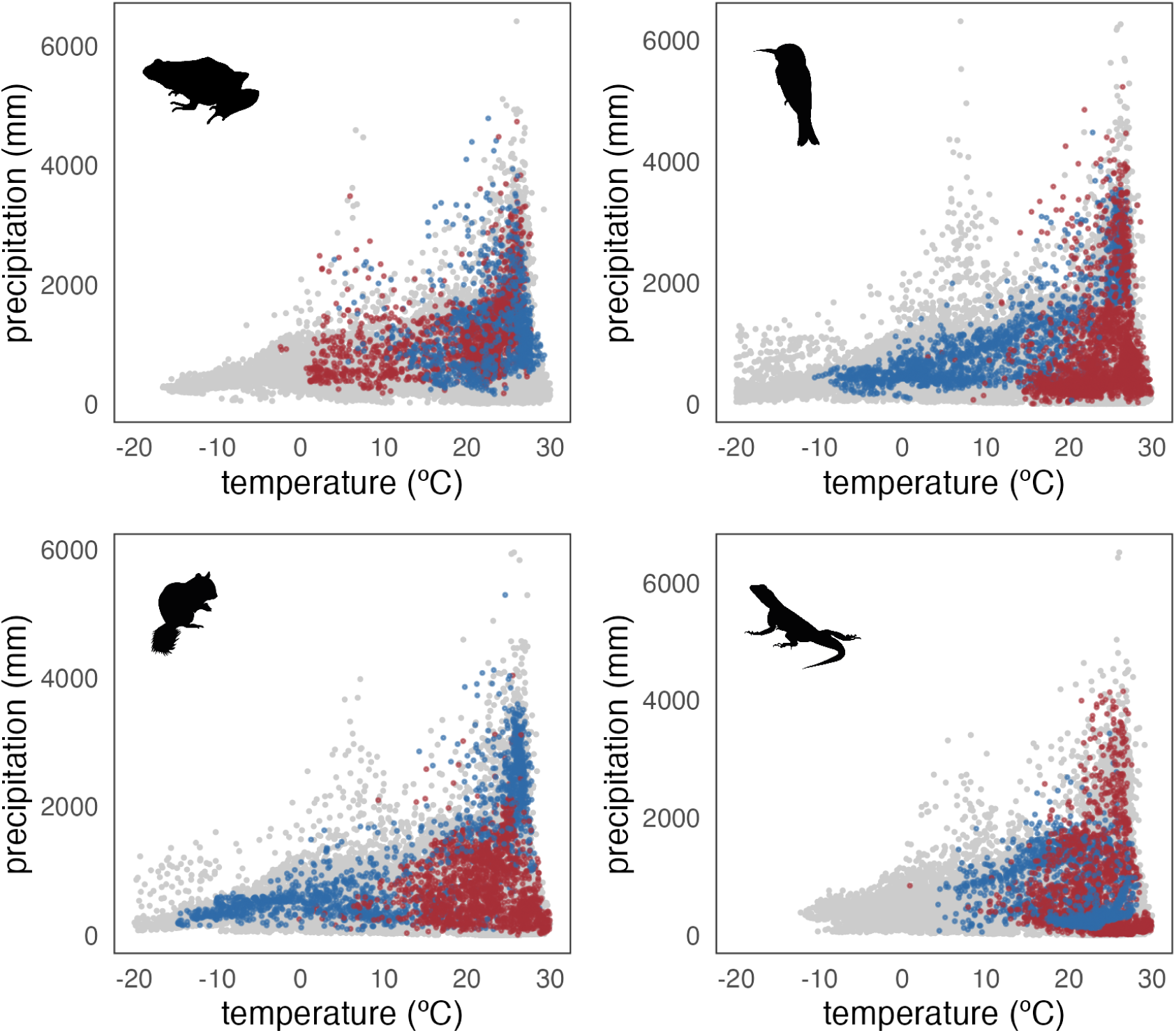
Climatic space (mean annual temperature vs. annual precipitation) occupied by regions of 10% highest (in red) and lowest (in blue) residual PD for each vertebrate clade. The grey points are grid cells with residual PD values between the 10th and 90th percentiles.

The patterns of distribution of regions with high and low residual PD across the environmental space defined by precipitation seasonality and temperature seasonality, or by NPP and TRI, do not follow clearly interpretable trends. There is no apparent segregation between high and low residual PD regions (Supp. Figures 16 and 17), except perhaps for birds in the climate seasonality space, where high residual PD regions tend to be at lower temperature seasonality than regions of low residual PD (Supp. Fig. 16).

Likewise, there is no apparent latitudinal gradient in residual PD for any vertebrate clade, in contrast to species richness (Supp. Fig. 18). Notably, regions of high richness for birds and to some extent for mammals across different latitudinal bands coincide with low residual PD levels (Supp. Fig. 18), although there appears to be no clear relationship between richness and residual PD (which is expected given that richness and PD are highly correlated).

## Discussion

In this study, we addressed the global geographic distribution of richness-corrected phylogenetic diversity (residual PD) for four major vertebrate clades (amphibians, birds, mammals, and squamates), as well as the effect of factors that potentially play a role in shaping large-scale diversity patterns.

The characterization of geographic patterns of residual PD reveal that, for all clades, continental Earth is mainly composed of large regions where either high or low levels of residual PD predominate (rather than a mixed or mosaic pattern of map cells with high and low residual PD that would be the expected outcome should the process generating the residual PD distribution be stochastic). We identified the areas of greatest concentration of high and low residual PD and found entire regions with consistently low (more closely related species than predicted by their species assemblage richness) and high (more distantly related species than predicted) levels for all four tetrapod groups (Fig. 2). Furthermore, we found a slight negative relationship of recent speciation rates with residual PD (Fig. 3), suggesting that recent diversification dynamics helped generate present-day global biogeographic patterns. Finally, our results show that evolutionary time (i.e., clade/lineage age) as well as most climatic variables had low to no effect on the differentiation of regions with highest and lowest residual PD (Figs. 4 and 5). Nonetheless, we identified differences between endotherms (birds and mammals) and ectotherms (amphibians and squamates) when considering temperature and precipitation levels in regions with highest and lowest values of residual PD (Fig. 5). This result indicates that thermal physiology may have influenced global diversity patterns among tetrapods.

We found that both American continents are regions of consistently low residual PD, meaning that regional assemblages are composed of species that are more closely related than expected by their richness. This applies to all the studied clades, with the exception of amphibians in North America and mammals in Eastern North America. In contrast, Africa consistently shows high levels of residual PD for all clades; in other words, it harbors species that are more distantly related than expected for all living tetrapods. Previous work on mammals (Davies and Buckley 2011) suggested that high residual PD found in Africa may reflect an African origin for many extant mammal clades (Lillegraven et al. 1987). While plausible for mammals, this hypothesis does not explain this pervasive biogeographic patterning in Africa across other major terrestrial vertebrate clades that likely originated on different continents (e.g., Benson et al. 2013; Claramunt and Cracraft 2015), and instead may be indicative of other shared factors related to the environmental and geological history of the African continent. Similarly, the low residual PD for all groups observed in America, particularly in South America (Fig. 2), may have been influenced by recent diversification events in multiple extant clades, particularly following dispersal after the formation of the Isthmus of Panama (ca. 3 million years ago; O’Dea et al. 2016), and coupled with the extinction of ancient endemic lineages (Webb 1976; Weir et al. 2009; Davies and Buckley 2011; Carrillo et al. 2020). This is also consistent with the ubiquitous negative relationship observed between residual PD and recent speciation rates which, although weak, reflects that recent speciation is likely one of the processes governing the geographic dimension of phylogenetic dynamics in vertebrates (Fig. 3). Taken together, these results may be indicative of recent climatic or geographic events (e.g., reconfiguration of continents) as primary drivers of recent speciation rates that have consequently shaped spatial patterns of extant tetrapod diversity.

In addition to abiotic processes influencing the spatial distribution of diversity, biotic factors likely drove some of these observable patterns (Davies and Buckley 2011). In particular, heterogeneous geographic residual PD patterns among clades (e.g., in Eurasia and Australia; Fig. 2) may indicate that intrinsic biological characteristics (i.e., physiological, ecological, or morphological) have played a role in the generation of such clade-specific patterns. For example, the Malay Archipelago shows particularly high residual PD for birds relative to other vertebrates, whereas parts of Australia present low residual PD for amphibians and squamates but high residual PD for mammals and birds (Fig. 2). It is plausible that these differences have arisen due to ecological and functional differences among clades, which may have determined the specific effects of climatic trends or tectonic processes in their biogeographic and evolutionary history. The high vagility of birds likely contributed to higher residual PD values in Southeast Asia and Australia, allowing repeated colonizations of islands by distantly related lineages in these regions (Jønsson and Fjeldså 2006; Sheldon et al. 2015). In addition, the high dispersal ability of most birds’ main subclades resulted in the arrival and relictual permanence of lineages with widely different phylogenetic origins (Jønsson et al. 2007), as opposed to isolated radiations (and, therefore, lower PD), which may be more frequent in organisms with lower dispersal abilities (Inger and Voris 2001; Siler et al. 2012). High residual PD of squamates within Southeast Asia may be related to multiple waves of island colonization during intervals of low sea level and environmental change (How and Kitchener 1997; Voris 2000; Brown et al. 2013; Husson et al. 2020), although this also affected mammals (Van Den Bergh et al. 2001; Meijaard 2003; Mercer and Roth 2003) and amphibians (Brown et al. 2013), which do not show comparable patterns of residual PD (perhaps due to a greater effect of environmental fluctuations promoting turnover in these taxa; e.g., Wilting et al. 2012).

In contrast, the low residual PD of squamates in arid Australia, and of amphibians in the temperate woodlands of the northern and eastern coasts of Australia, likely resulted from a few extensive radiations (see the phylogenetic trees in Fig. 2). These radiations may be facilitated by adaptation and specialization within these unique biomes, leading to community assembly driven by proportionally more closely related species (Pianka 1981; Rabosky et al. 2007; Skinner et al. 2011; Vidal-García and Keogh 2015; Tejero-Cicuéndez et al. 2022; Brennan et al. 2023). Finally, geographic patterns of residual PD may be partially generated by ancient evolutionary lineages that inhabit certain regions. This could be the case of dibamid reptiles in Southeast Asia (Townsend et al. 2011), palaeognath birds in Australia (Yonezawa et al. 2017), or marsupial and xenarthran mammals in Australia and North America, respectively (May-Collado et al. 2015). Additionally, passerine birds, that comprise ∼60% of extant avian diversity, contribute to high residual PD in Australia due to the presence of numerous endemic lineages that are distantly related in comparison to the more diverse, but generally closely related, passerine lineages inhabiting South America (Moyle et al. 2016; Oliveros et al. 2019; Harvey et al. 2020; Machac 2020) (Fig. 2). These contrasting patterns are somewhat paradoxical considering that passerines are generally competent fliers, suggesting that there may be additional factors driving spatial diversity patterns among birds.

Previous work has identified that diversity dynamics are strongly influenced by both extrinsic (e.g., the paleogeographic history) and intrinsic (e.g., ecomorphological or niche-related) factors (James and Shine 2000; Jetz and Rahbek 2002; Badgley 2010; Pyron 2014a; Menéndez et al. 2021; Jiang et al. 2023). For instance, mountainous regions (a universally recognized driver of diversity and evolutionary processes; Hoorn et al. 2010; Rahbek et al. 2019; Perrigo et al. 2020) serve multiple functions during biodiversity generation; mountains can promote speciation due to habitat heterogeneity across an elevational gradient (acting as a source of diversity, particularly in the tropics; Cadena et al. 2011; Šmíd et al. 2021), or induce dispersal barriers (e.g., Miller et al. 2008). Mountains may also generate important refugia and reserves of cold- or humid-adapted diversity during periods of climate warming and aridification (Hampe and Jump 2011; Fjeldså et al. 2012). Likewise, deserts, which have been recurrently considered sinks of diversity (i.e., regions harboring distantly related species due to a lack of within-system diversification; Crisp et al. 2009) may in fact harbor low levels of residual PD, as our results show in Australia for squamates and North America for birds, mammals and squamates (Fig. 2). In many cases, this may reflect large radiations of certain clades specialized to the arid conditions (e.g., Wiens et al. 2013; Rabosky et al. 2014). This ecological versatility of physiographic features may be responsible for the apparent decoupling between environmental variables and residual PD (Supp. Figs.10-15). However, we found some segregation in the climate space between the regions of highest and lowest residual PD, especially in birds and mammals (Fig. 5). This may be explained by fundamental physiological differences between endotherms and ectotherms (Buckley et al. 2012), which presumably determine the impact of climatic events on their diversity patterns.

Apart from differences in trait- and environment-mediated speciation and dispersal, extinction may also be a major driver of diversity, diversification and biogeographic patterns (Jablonski 2001; Seeholzer and Brumfield 2023). Specifically, extinction events are known not only to underlie current patterns of species richness (Pyron 2014b; Meseguer and Condamine 2020), but also to substantially affect other facets of biodiversity (Erwin 2008; Pimiento et al. 2020; Brocklehurst et al. 2021). Extinction may both increase and decrease phylogenetic diversity, depending on the age and phylogenetic clustering of the lineages that are more prone to extinction (Daru et al. 2017). High residual PD regions may result from higher extinction rates of species from relatively recent radiations, mainly reducing species assemblages to taxa with more distant evolutionary relationships, whereas low residual PD may arise in regions where extinction rates are higher for relatively old diversity (Vasconcelos et al. 2022). The exploration of extinction dynamics and, critically, the inclusion of fossil data (which enables better estimates of extinction and deeper speciation events; Quintero et al. 2023), will help to further disentangle the evolutionary factors underpinning geographic patterns of vertebrate diversity. Likewise, our results show no apparent relationship between the residual PD of a region and the age of the biota within it or the pattern of lineage accumulation (Supp. Figs. 5-9), but the inclusion of fossil information would enhance our ability to investigate such a relationship.

### Conservation implications

Our study provides a global perspective of diversity dynamics in a comparative framework, which allows us to explore the processes that have shaped the heterogeneous evolutionary history of tetrapods. Recent speciation rates are responsible for a significant part of the patterns we report, but there are other factors likely implicated, such as the differential effects of climate on endotherms and ectotherms due to specialized adaptations, deep-time speciation dynamics, and extinction events. Future studies focused on diversification-through-time patterns and on the mechanistic aspects of adaptation to different environmental conditions (e.g., physiology, morphology) are needed to further elucidate the factors at play in global biodiversity patterns. Moving forward, our methodological approach to explore spatial phylogenetic diversity and its drivers may help to inform conservation policies beyond the species-richness-hotspot and endemism strategies (Myers et al. 2000). Integrating evolutionary relationships at a regional scale has been stated as a necessary step for global conservation efforts for decades (Faith 1992), but the direct comparison of different clades may be critical to identify biodiversity priorities. Assessing patterns of residual PD for focal clades would inform on which regions harbor the highest lineage diversity (highest residual PD), facilitating efforts to maximize conservation of phylogenetically distinct lineages and help to preserve larger portions of the evolutionary history of entire clades. Additionally, regions with the lowest residual PD that may be acting as sources of diversity (Meseguer et al. 2020) can be considered in conservation initiatives as priority areas in order to safeguard the generation of new biodiversity. Furthermore, integrating paleontological data into our method would facilitate a novel, multidisciplinary approach for utilizing deep time data for species conservation prioritization (Pimiento and Antonelli 2022).

## Competing interests

The authors declare no competing interests.

## Notes

### Competing Interest Statement

The authors have declared no competing interest.

### Summary of Updates

Objectives and hypotheses stated more explicitly, and temporal patterns analyzed more rigorously. Text revised to improve readability. Figures updated.

### Funding

- Agencia Estatal de InvestigaciM_CM_3n: 10.13039/501100011033
- Alexander von Humboldt Foundation
- UK Research and Innovation: MR/X015130/1
- Swedish Research Council: 2019-04739
- Czech Science Foundation: 22-12757S
- Charles University: 204069

## References

Allee W.C. 1926. Distribution of Animals in a Tropical Rain-Forest with Relation to Environmental Factors. Ecology. 7:445–468.

Anderson S. 1974. Patterns of Faunal Evolution. The Quarterly Review of Biology. 49:311–332.

Badgley C. 2010. Tectonics, topography, and mammalian diversity. Ecography. 33:220–231.

Benson R.B.J., Mannion P.D., Butler R.J., Upchurch P., Goswami A., Evans S.E. 2013. Cretaceous tetrapod fossil record sampling and faunal turnover: Implications for biogeography and the rise of modern clades. Palaeogeography, Palaeoclimatology, Palaeoecology. 372:88–107.

Brennan I.G., Lemmon A.R., Lemmon E.M., Hoskin C.J., Donnellan S.C., Keogh J.S. 2023. Populating a Continent: Phylogenomics Reveal the Timing of Australian Frog Diversification. Systematic Biology.:syad048.

Brocklehurst N., Panciroli E., Benevento G.L., Benson R.B.J. 2021. Mammaliaform extinctions as a driver of the morphological radiation of Cenozoic mammals. Current Biology. 31:2955–2963.e4.

Brown R.M., Siler C.D., Oliveros C.H., Esselstyn J.A., Diesmos A.C., Hosner P.A., Linkem C.W., Barley A.J., Oaks J.R., Sanguila M.B., Welton L.J., Blackburn D.C., Moyle R.G., Townsend Peterson A., Alcala A.C. 2013. Evolutionary Processes of Diversification in a Model Island Archipelago. Annual Review of Ecology, Evolution, and Systematics. 44:411–435.

Buckley L.B., Hurlbert A.H., Jetz W. 2012. Broad-scale ecological implications of ectothermy and endothermy in changing environments. Global Ecology and Biogeography. 21:873–885.

Cadena C.D., Kozak K.H., Gómez J.P., Parra J.L., McCain C.M., Bowie R.C.K., Carnaval A.C., Moritz C., Rahbek C., Roberts T.E., Sanders N.J., Schneider C.J., VanDerWal J., Zamudio K.R., Graham C.H. 2011. Latitude, elevational climatic zonation and speciation in New World vertebrates. Proceedings of the Royal Society B: Biological Sciences. 279:194–201.

de Candolle A.L.P.P. 1859. On the Causes which Limit Vegetable Species Towards the North, in Europe and Similar Regions. Annual Report of the Board of Regents of the Smithsonian Institution for the Year 1858.:237–245.

Carrillo J.D., Faurby S., Silvestro D., Jaramillo C., Bacon C.D., Antonelli A. 2020. Disproportionate extinction of South American mammals drove the asymmetry of the Great American Biotic Interchange. Proceedings of the National Academy of Sciences.:1–7.

Claramunt S., Cracraft J. 2015. A new time tree reveals Earth history’s imprint on the evolution of modern birds. Sci. Adv. 1:e1501005.

Collyer M.L., Adams D.C. 2018. RRPP: An r package for fitting linear models to high-dimensional data using residual randomization. Methods in Ecology and Evolution. 9:1772–1779.

Collyer M.L., Adams D.C. 2022. RRPP: Linear Model Evaluation with Randomized Residuals in a Permutation Procedure..

Crisp M.D., Arroyo M.T.K., Cook L.G., Gandolfo M.A., Jordan G.J., McGlone M.S., Weston P.H., Westoby M., Wilf P., Linder H.P. 2009. Phylogenetic biome conservatism on a global scale. Nature. 458:754–756.

Daru B.H., Elliott T.L., Park D.S., Davies T.J. 2017. Understanding the Processes Underpinning Patterns of Phylogenetic Regionalization. Trends in Ecology & Evolution. 32:845–860.

Darwin C. 1859. On the Origin of Species by Means of Natural Selection, or the Preservation of Favoured Races in the Struggle for Life. London, UK: John Murray.

Davies T.J., Buckley L.B. 2011. Phylogenetic diversity as a window into the evolutionary and biogeographic histories of present-day richness gradients for mammals. Philosophical Transactions of the Royal Society B: Biological Sciences. 366:2414–2425.

Dobzhansky T. 1950. Evolution in the tropics. American Scientist. 38:209–221.

Earl C., Belitz M.W., Laffan S.W., Barve V., Barve N., Soltis D.E., Allen J.M., Soltis P.S., Mishler B.D., Kawahara A.Y., Guralnick R. 2021. Spatial phylogenetics of butterflies in relation to environmental drivers and angiosperm diversity across North America. iScience. 24:102239.

Erwin D.H. 2008. Extinction as the loss of evolutionary history. Proceedings of the National Academy of Sciences. 105:11520–11527.

Faith D.P. 1992. Conservation evaluation and phylogenetic diversity. Biological Conservation. 61:1–10.

Fick S.E., Hijmans R.J. 2017. WorldClim 2: new 1-km spatial resolution climate surfaces for global land areas. International Journal of Climatology. 37:4302–4315.

Fischer A.G. 1960. Latitudinal variations in organic diversity. Evolution. 14:64–81.

Fjeldså J., Bowie R.C.K., Rahbek C. 2012. The Role of Mountain Ranges in the Diversification of Birds. Annual Review of Ecology, Evolution, and Systematics. 43:249–265.

Fritz S.A., Rahbek C. 2012. Global patterns of amphibian phylogenetic diversity. Journal of biogeography. 39:1373–1382.

Gumbs R., Gray C.L., Böhm M., Hoffmann M., Grenyer R., Jetz W., Meiri S., Roll U., Owen N.R., Rosindell J. 2020. Global priorities for conservation of reptilian phylogenetic diversity in the face of human impacts. Nat Commun. 11:2616.

Hackett S.J., Kimball R.T., Reddy S., Bowie R.C.K., Braun E.L., Braun M.J., Chojnowski J.L., Cox W.A., Han K.-L., Harshman J., Huddleston C.J., Marks B.D., Miglia K.J., Moore W.S., Sheldon F.H., Steadman D.W., Witt C.C., Yuri T. 2008. A Phylogenomic Study of Birds Reveals Their Evolutionary History. Science. 320:1763–1768.

Hampe A., Jump A.S. 2011. Climate Relicts: Past, Present, Future. Annual Review of Ecology, Evolution, and Systematics. 42:313–333.

Harmon L.J. 2012. An Inordinate Fondness for Eukaryotic Diversity. PLoS Biology. 10:8–11.

Harvey M.G., Bravo G.A., Claramunt S., Cuervo A.M., Derryberry G.E., Battilana J., Seeholzer G.F., McKay J., O’Meara B.C., Faircloth B.C., Edwards S.V., Pérez-Emán J.L., Moyle R.G., Sheldon F.H., Aleixo A., Brian Tilston Smith, Smith B.D., Chesser R.T., Silveira L.F., Cracraft J., Brumfield R.T., Derryberry E.P. 2020. The evolution of a tropical biodiversity hotspot. Science. 370:1343–1348.

Hoorn C., Wesselingh F.P., ter Steege H., Bermudez M.A., Mora A., Sevink J., Sanmartín I., Sanchez-Meseguer A., Anderson C.L., Figueiredo J.P., Jaramillo C., Riff D., Negri F.R., Hooghiemstra H., Lundberg J., Stadler T., Särkinen T., Antonelli A. 2010. Amazonia Through Time: Andean Uplift, Climate Change, Landscape Evolution, and Biodiversity. Science. 330:927–931.

How R.A., Kitchener D.J. 1997. Biogeography of Indonesian snakes. Journal of Biogeography. 24:725–735.

von Humboldt A., Bonpland A. 1807. Essai sur la géographie des plantes. Paris, France: Schoell.

Husson L., Boucher F.C., Sarr A.-C., Sepulchre P., Cahyarini S.Y. 2020. Evidence of Sundaland’s subsidence requires revisiting its biogeography. Journal of Biogeography. 47:843–853.

Inger R.F., Voris H.K. 2001. The biogeographical relations of the frogs and snakes of Sundaland. Journal of Biogeography. 28:863–891.

IUCN. 2022. The IUCN Red List of Threatened Species. Version 2022–2. Available from https://www.iucnredlist.org.

Jablonski D. 2001. Lessons from the past: Evolutionary impacts of mass extinctions. Proceedings of the National Academy of Sciences. 98:5393–5398.

James C.D., Shine R. 2000. Why are there so many coexisting species of lizards in Australian deserts? Oecologia. 125:127–141.

Jetz W., Pyron R.A. 2018. The interplay of past diversification and evolutionary isolation with present imperilment across the amphibian tree of life. Nature Ecology and Evolution. 2:850–858.

Jetz W., Rahbek C. 2002. Geographic range size and determinants of avian species richness. Science. 297:1548–1551.

Jetz W., Thomas G.H., Joy J.B., Hartmann K., Mooers A.O. 2012. The global diversity of birds in space and time. Nature. 491:444–448.

Jiang K., Wang Q., Dimitrov D., Luo A., Xu X., Su X., Liu Y., Li Y., Li Y., Wang Z. 2023. Evolutionary history and global angiosperm species richness–climate relationships. Global Ecology and Biogeography. 32:1059–1072.

Jønsson K.A., Fjeldså J. 2006. Determining biogeographical patterns of dispersal and diversification in oscine passerine birds in Australia, Southeast Asia and Africa. Journal of Biogeography. 33:1155–1165.

Jønsson K.A., Fjeldså J., Ericson P.G.P., Irestedt M. 2007. Systematic placement of an enigmatic Southeast Asian taxon *Eupetes macrocerus* and implications for the biogeography of a main songbird radiation, the Passerida. Biology Letters. 3:323–326.

Lillegraven J.A., Thompson S.D., McNab B.K., Patton J.L. 1987. The origin of eutherian mammals. Biological Journal of the Linnean Society. 32:281–336.

Louca S., Pennell M.W. 2020. Extant timetrees are consistent with a myriad of diversification histories. Nature. 580:502–505.

MacArthur R.H. 1965. Patterns of Species Diversity. Biological Reviews. 40:510–533.

Machac A. 2020. The dynamics of bird diversity in the new world. Systematic Biology. 69:1180–1199.

Matthew W.D. 1915. Climate and evolution. Annals of the New York Academy of Sciences. 24:171–318.

May-Collado L.J., Kilpatrick C.W., Agnarsson I. 2015. Mammals from ‘down under’: a multi-gene species-level phylogeny of marsupial mammals (Mammalia, Metatheria). PeerJ. 3:e805.

Meijaard E. 2003. Mammals of south-east Asian islands and their Late Pleistocene environments. Journal of Biogeography. 30:1245–1257.

Menéndez I., Gómez Cano A.R., Cantalapiedra J.L., Peláez-Campomanes P., Álvarez-Sierra M.Á., Hernández Fernández M. 2021. A multi-layered approach to the diversification of squirrels. Mammal Review. 51:66–81.

Mercer J.M., Roth V.L. 2003. The Effects of Cenozoic Global Change on Squirrel Phylogeny. Science. 299:1568–1572.

Meseguer A.S., Antoine P.-O., Fouquet A., Delsuc F., Condamine F.L. 2020. The role of the Neotropics as a source of world tetrapod biodiversity. Global Ecology and Biogeography. 29:1565–1578.

Meseguer A.S., Condamine F.L. 2020. Ancient tropical extinctions at high latitudes contributed to the latitudinal diversity gradient. Evolution. 74:1966–1987.

Miller M.J., Bermingham E., Klicka J., Escalante P., do Amaral F.S.R., Weir J.T., Winker K. 2008. Out of Amazonia again and again: episodic crossing of the Andes promotes diversification in a lowland forest flycatcher. Proceedings of the Royal Society B: Biological Sciences. 275:1133–1142.

Mishler B.D. 2023. Spatial phylogenetics. Journal of Biogeography. 50:1454–1463.

Mittelbach G.G., Schemske D.W., Cornell H.V., Allen A.P., Brown J.M., Bush M.B., Harrison S.P., Hurlbert A.H., Knowlton N., Lessios H.A., McCain C.M., McCune A.R., McDade L.A., McPeek M.A., Near T.J., Price T.D., Ricklefs R.E., Roy K., Sax D.F., Schluter D., Sobel J.M., Turelli M. 2007. Evolution and the latitudinal diversity gradient: Speciation, extinction and biogeography. Ecology Letters. 10:315–331.

Moore B. 1920. The Ecological Society and Its Opportunity. Science. 51:67–68.

Morlon H., Schwilk D.W., Bryant J.A., Marquet P.A., Rebelo A.G., Tauss C., Bohannan B.J.M., Green J.L. 2011. Spatial patterns of phylogenetic diversity. Ecology Letters. 14:141–149.

Moyle R.G., Oliveros C.H., Andersen M.J., Hosner P.A., Benz B.W., Manthey J.D., Travers S.L., Brown R.M., Faircloth B.C. 2016. Tectonic collision and uplift of Wallacea triggered the global songbird radiation. Nature Communications. 7:12709.

Myers N., Mittermeier R.A., Mittermeier C.G., da Fonseca G.A.B., Kent J. 2000. Biodiversity hotspots for conservation priorities. Nature. 403:853–858.

Nee S., May R.M. 1997. Extinction and the Loss of Evolutionary History. Science. 278:692–694.

O’Dea A., Lessios H.A., Coates A.G., Eytan R.I., Restrepo-Moreno S.A., Cione A.L., Collins L.S., de Queiroz A., Farris D.W., Norris R.D., Stallard R.F., Woodburne M.O., Aguilera O., Aubry M.-P., Berggren W.A., Budd A.F., Cozzuol M.A., Coppard S.E., Duque-Caro H., Finnegan S., Gasparini G.M., Grossman E.L., Johnson K.G., Keigwin L.D., Knowlton N., Leigh E.G., Leonard-Pingel J.S., Marko P.B., Pyenson N.D., Rachello-Dolmen P.G., Soibelzon E., Soibelzon L., Todd J.A., Vermeij G.J., Jackson J.B.C. 2016. Formation of the Isthmus of Panama. Science Advances. 2:e1600883.

Oliveros C.H., Field D.J., Ksepka D.T., Barker F.K., Aleixo A., Andersen M.J., Alström P., Benz B.W., Braun E.L., Braun M.J., Bravo G.A., Brumfield R.T., Chesser R.T., Claramunt S., Cracraft J., Cuervo A.M., Derryberry E.P., Glenn T.C., Harvey M.G., Hosner P.A., Joseph L., Kimball R.T., Mack A.L., Miskelly C.M., Peterson A.T., Robbins M.B., Sheldon F.H., Silveira L.F., Smith B.T., White N.D., Moyle R.G., Faircloth B.C. 2019. Earth history and the passerine superradiation. Proceedings of the National Academy of Sciences.

Paradis E., Schliep K. 2019. ape 5.0: an environment for modern phylogenetics and evolutionary analyses in R. Bioinformatics. 35:526–528.

Pennell M.W., Eastman J.M., Slater G.J., Brown J.W., Uyeda J.C., Fitzjohn R.G., Alfaro M.E., Harmon L.J. 2014. Geiger v2.0: An expanded suite of methods for fitting macroevolutionary models to phylogenetic trees. Bioinformatics. 30:2216–2218.

Perrigo A., Hoorn C., Antonelli A. 2020. Why mountains matter for biodiversity. Journal of Biogeography.:315–325.

Pianka E.R. 1981. Diversity and adaptive radiations of Australian desert lizards. Ecological biogeography of Australia. p. 1376–1392.

Pimiento C., Antonelli A. 2022. Integrating deep-time palaeontology in conservation prioritisation. Frontiers in Ecology and Evolution. 10.

Pimiento C., Bacon C.D., Silvestro D., Hendy A., Jaramillo C., Zizka A., Meyer X., Antonelli A., Bacon C.D. 2020. Selective extinction against redundant species buffers functional diversity. Proceedings of the Royal Society B: Biological Sciences. 287:20201162.

Pinzon J.E., Tucker C.J. 2014. A non-stationary 1981-2012 AVHRR NDVI3g time series. Remote Sensing. 6:6929–6960.

Pybus O.G., Harvey P.H. 2000. Testing macro–evolutionary models using incomplete molecular phylogenies. Proceedings of the Royal Society of London. Series B: Biological Sciences. 267:2267–2272.

Pyron R.A. 2014a. Biogeographic analysis reveals ancient continental vicariance and recent oceanic dispersal in amphibians. Systematic Biology. 63:779–797.

Pyron R.A. 2014b. Temperate extinction in squamate reptiles and the roots of latitudinal diversity gradients. Global Ecology and Biogeography. 23:1126–1134.

Pyron R.A., Wiens J.J. 2013. Large-scale phylogenetic analyses reveal the causes of high tropical amphibian diversity. Proceedings of the Royal Society B: Biological Sciences. 280:20131622.

Quintero I., Landis M.J., Jetz W., Morlon H. 2023. The build-up of the present-day tropical diversity of tetrapods. Proceedings of the National Academy of Sciences. 120:e2220672120.

R Core Team. 2023. R: A Language and Environment for Statistical Computing. Vienna, Austria: R Foundation for Statistical Computing.

Rabosky D.L., Donnellan S.C., Talaba A.L., Lovette I.J. 2007. Exceptional among-lineage variation in diversification rates during the radiation of Australia’s most diverse vertebrate clade. Proceedings of the Royal Society B: Biological Sciences. 274:2915–2923.

Rabosky D.L., Hutchinson M.N., Donnellan S.C., Talaba A.L., Lovette I.J. 2014. Phylogenetic disassembly of species boundaries in a widespread group of Australian skinks (Scincidae: *Ctenotus*). Molecular Phylogenetics and Evolution. 77:71–82.

Rahbek C., Borregaard M.K., Antonelli A., Colwell R.K., Holt B.G., Nogues-Bravo D., Rasmussen C.M.Ø., Richardson K., Rosing M.T., Whittaker R.J., Fjeldså J. 2019. Building mountain biodiversity: Geological and evolutionary processes. Science. 365:1114–1119.

Redding D.W., Mooers A.O. 2006. Incorporating evolutionary measures into conservation prioritization. Conservation Biology. 20:1670–1678.

Ren K. 2021. rlist: A toolbox for non-tabular data manipulation.

Revell L.J. 2012. phytools: An R package for phylogenetic comparative biology (and other things). Methods in Ecology and Evolution. 3:217–223.

Ricklefs R.E. 2004. A comprehensive framework for global patterns in biodiversity. Ecology Letters. 7:1–15.

Rodrigues A.S.L., Brooks T.M., Gaston K.J. 2005. Integrating phylogenetic diversity in the selection of priority areas for conservation: does it make a difference? In: Purvis A., Gittleman J.L., Brooks T., editors. Phylogeny and Conservation. Cambridge University Press.

Roll U., Meiri S., Farrell M., Davies J., Gittleman J., Wiens J., Stephens P. 2021. GARD 1.5 range shapefiles used in: Global diversity patterns are explained by diversification rates at ancient, not shallow, timescales..

Ruthven A.G. 1920. The Environmental Factors in the Distribution of Animals. Geographical Review. 10:241–248.

Saupe E.E. 2023. Explanations for latitudinal diversity gradients must invoke rate variation. Proceedings of the National Academy of Sciences. 120:e2306220120.

Seeholzer G.F., Brumfield R.T. 2023. Speciation-by-Extinction. Systematic Biology.:syad049.

Sheldon F.H., Lim H.C., Moyle R.G. 2015. Return to the Malay Archipelago: the biogeography of Sundaic rainforest birds. J Ornithol. 156:91–113.

Siler C.D., Oaks J.R., Welton L.J., Linkem C.W., Swab J.C., Diesmos A.C., Brown R.M. 2012. Did geckos ride the Palawan raft to the Philippines? Journal of Biogeography. 39:1217–1234.

Skinner A., Hugall A.F., Hutchinson M.N. 2011. Lygosomine phylogeny and the origins of Australian scincid lizards. Journal of Biogeography. 38:1044–1058.

Šmíd J., Sindaco R., Shobrak M., Busais S., Tamar K., Aghová T., Simó-Riudalbas M., Tarroso P., Geniez P., Crochet P.A., Els J., Burriel-Carranza B., Tejero-Cicuéndez H., Carranza S. 2021. Diversity patterns and evolutionary history of Arabian squamates. Journal of Biogeography. 48:1183–1199.

Stebbins G.L. 1974. Flowering plants: evolution above the species level. Harvard University Press.

Tejero-Cicuéndez H., Tarroso P., Carranza S., Rabosky D.L. 2022. Desert lizard diversity worldwide: Effects of environment, time, and evolutionary rate. Global Ecology and Biogeography. 31:776–790.

Title P.O., Bemmels J.B. 2018. ENVIREM: an expanded set of bioclimatic and topographic variables increases flexibility and improves performance of ecological niche modeling. Ecography. 41:291–307.

Title P.O., Daniel L. Rabosky, Daniel L. Rabosky, Daniel L. Rabosky, Rabosky D.L. 2019. Tip rates, phylogenies and diversification: What are we estimating, and how good are the estimates? Methods in Ecology and Evolution. 10:821–834.

Title P.O., Swiderski D.L., Zelditch M.L. 2022. EcoPhyloMapper: An r package for integrating geographical ranges, phylogeny and morphology. Methods in Ecology and Evolution. 13:1912–1922.

Tonini J.F.R., Beard K.H., Ferreira R.B., Jetz W., Pyron R.A. 2016. Fully-sampled phylogenies of squamates reveal evolutionary patterns in threat status. Biological Conservation. 204:23–31.

Townsend T.M., Leavitt D.H., Reeder T.W. 2011. Intercontinental dispersal by a microendemic burrowing reptile (Dibamidae). Proceedings of the Royal Society B: Biological Sciences. 278:2568–2574.

Tucker C.M., Cadotte M.W. 2013. Unifying measures of biodiversity: understanding when richness and phylogenetic diversity should be congruent. Diversity and Distributions. 19:845–854.

Tucker C.M., Cadotte M.W., Carvalho S.B., Davies T.J., Ferrier S., Fritz S.A., Grenyer R., Helmus M.R., Jin L.S., Mooers A.O., Pavoine S., Purschke O., Redding D.W., Rosauer D.F., Winter M., Mazel F. 2017. A guide to phylogenetic metrics for conservation, community ecology and macroecology. Biological Reviews. 92:698–715.

Upham N.S., Esselstyn J.A., Jetz W. 2019. Inferring the mammal tree: Species-level sets of phylogenies for questions in ecology, evolution, and conservation. PLoS Biology. 17:1–44.

Van Den Bergh G.D., De Vos J., Sondaar P.Y. 2001. The Late Quaternary palaeogeography of mammal evolution in the Indonesian Archipelago. Palaeogeography, Palaeoclimatology, Palaeoecology. 171:385–408.

Vasconcelos T., O’Meara B.C., Beaulieu J.M. 2022. Retiring “Cradles” and “Museums” of Biodiversity. The American Naturalist. 199:195–204.

Vásquez-Restrepo J.D., Ochoa-Ochoa L.M., Flores-Villela O., Velasco J.A. 2023. Deconstructing the dimensions of alpha diversity in squamate reptiles (Reptilia: Squamata) across the Americas. Global Ecology and Biogeography. 32:250–266.

Velasco J.A., Pinto-Ledezma J.N. 2022. Mapping species diversification metrics in macroecology: Prospects and challenges. Frontiers in Ecology and Evolution. 10:951271.

Vidal-García M., Keogh J.S. 2015. Convergent evolution across the Australian continent: ecotype diversification drives morphological convergence in two distantly related clades of Australian frogs. Journal of Evolutionary Biology. 28:2136–2151.

Voris H.K. 2000. Maps of Pleistocene sea levels in Southeast Asia: shorelines, river systems and time durations. Journal of Biogeography. 27:1153–1167.

Voskamp A., Baker D.J., Stephens P.A., Valdes P.J., Willis S.G. 2017. Global patterns in the divergence between phylogenetic diversity and species richness in terrestrial birds. Journal of Biogeography. 44:709–721.

Wallace A.R. 1876. The geographical distribution of animals. New York, USA: Harper and Brothers.

Webb D.S. 1976. Mammalian faunal dynamics of the great American interchange. Paleobiology. 2:220–234.

Weir J.T., Bermingham E., Schluter D. 2009. The Great American Biotic Interchange in birds. Proceedings of the National Academy of Sciences. 106:21737–21742.

Whittaker R.H. 1975. Communities and ecosystems. Macmillan Publishing.

Wiens J.J. 2011. The causes of species richness patterns across space, time, and clades and the role of “ecological limits.” Quarterly Review of Biology. 86:75–96.

Wiens J.J. 2015. Explaining large-scale patterns of vertebrate diversity. Biology Letters. 11:20150506.

Wiens J.J., Kozak K.H., Silva N. 2013. Diversity and niche evolution along aridity gradients in North American lizards (Phrynosomatidae). Evolution. 67:1715–1728.

Wilson M.F.J., O’Connell B., Brown C., Guinan J.C., Grehan A.J. 2007. Multiscale Terrain Analysis of Multibeam Bathymetry Data for Habitat Mapping on the Continental Slope. Marine Geodesy. 30:3–35.

Wilting A., Sollmann R., Meijaard E., Helgen K.M., Fickel J. 2012. Mentawai’s endemic, relictual fauna: is it evidence for Pleistocene extinctions on Sumatra? Journal of Biogeography. 39:1608–1620.

Yonezawa T., Segawa T., Mori H., Campos P.F., Hongoh Y., Endo H., Akiyoshi A., Kohno N., Nishida S., Wu J., Jin H., Adachi J., Kishino H., Kurokawa K., Nogi Y., Tanabe H., Mukoyama H., Yoshida K., Rasoamiaramanana A., Yamagishi S., Hayashi Y., Yoshida A., Koike H., Akishinonomiya F., Willerslev E., Hasegawa M. 2017. Phylogenomics and Morphology of Extinct Paleognaths Reveal the Origin and Evolution of the Ratites. Current Biology. 27:68–77.

