## Supplemental Tables for "The role of recent speciation in present-day patterns of tetrapod phylogenetic relatedness"

**Supplementary Table 1**. Results from the linear model of recent speciation (DR) rates against residual phylogenetic diversity (PD).

| **Clade** | **Estimate** | **R.squared** | **p.value** |
|---|---|---|---|
| **amphibians** | -2.09E-05 | 0.005 | 1.71E-16 |
| **birds** | -7.24E-05 | 0.047 | 6.82E-209 |
| **mammals** | -0.0001676 | 0.076 | 9.96E-274 |
| **squamates** | -5.74E-05 | 0.125 | 0 |

**Supplementary Table 2**. Post-hoc analyses comparing DR rates between pairs of focal regions of high and low residual PD.

| | taxa | d | UCL (95%) | Z | Pr > d |
|---|---|---|---|---|---|
| cradle_australia_oceania:cradle_oriental | amphibians | 0.00525528 | 0.00698079 | 1.10233047 | 0.15 |
| cradle_australia_oceania:cradle_south_america | amphibians | 0.01561991 | 0.00578026 | 3.57674856 | 0.001 |
| cradle_australia_oceania:museum_africa | amphibians | 0.0114298 | 0.0065061 | 2.74666907 | 0.003 |
| cradle_oriental:cradle_south_america | amphibians | 0.01036463 | 0.00647359 | 2.49136758 | 0.002 |
| cradle_oriental:museum_africa | amphibians | 0.01668508 | 0.00701332 | 3.55794931 | 0.001 |
| cradle_south_america:museum_africa | amphibians | 0.02704971 | 0.00531601 | 5.46148224 | 0.001 |
| cradle_north_america:cradle_south_america | birds | 0.00267518 | 0.01626041 | -0.7325484 | 0.755 |
| cradle_north_america:museum_africa_madagascar | birds | 0.04391713 | 0.01936177 | 3.34503922 | 0.001 |
| cradle_north_america:museum_australia_oceania_asia | birds | 0.05353809 | 0.01732115 | 4.1915594 | 0.001 |
| cradle_south_america:museum_africa_madagascar | birds | 0.04124195 | 0.01537753 | 3.70799933 | 0.001 |
| cradle_south_america:museum_australia_oceania_asia | birds | 0.05086291 | 0.01456124 | 4.46237966 | 0.001 |
| museum_africa_madagascar:museum_australia_oceania_asia | birds | 0.00962096 | 0.01660227 | 0.69577414 | 0.257 |
| cradle_north_america:cradle_south_america1 | mammals | 0.02791751 | 0.02450657 | 1.76948243 | 0.027 |
| cradle_north_america:museum_africa | mammals | 0.00283974 | 0.02563416 | -0.9689377 | 0.827 |
| cradle_north_america:museum_north_america | mammals | 0.02831498 | 0.03051384 | 1.43523423 | 0.072 |
| cradle_south_america:museum_africa1 | mammals | 0.03075725 | 0.01946118 | 2.5414738 | 0.004 |
| cradle_south_america:museum_north_america | mammals | 0.05623249 | 0.02699161 | 2.99347373 | 0.001 |
| museum_africa:museum_north_america | mammals | 0.02547524 | 0.02410014 | 1.6703083 | 0.044 |
| cradle_australia:cradle_north_america | squamates | 0.04805722 | 0.01229255 | 4.67906539 | 0.001 |
| cradle_australia:cradle_south_america | squamates | 0.09239165 | 0.00982233 | 7.93397381 | 0.001 |
| cradle_australia:museum_south_africa | squamates | 0.0168303 | 0.01091905 | 2.46286027 | 0.001 |
| cradle_australia:museum_southeast_asia | squamates | 0.01053019 | 0.0096155 | 1.74174676 | 0.029 |
| cradle_north_america:cradle_south_america2 | squamates | 0.04433443 | 0.01151994 | 4.8334128 | 0.001 |
| cradle_north_america:museum_south_africa | squamates | 0.06488752 | 0.01223027 | 6.1095102 | 0.001 |
| cradle_north_america:museum_southeast_asia | squamates | 0.03752703 | 0.01114818 | 4.40339537 | 0.001 |
| cradle_south_america:museum_south_africa | squamates | 0.10922195 | 0.01094116 | 8.42007702 | 0.001 |
| cradle_south_america:museum_southeast_asia | squamates | 0.08186147 | 0.00954673 | 7.67088747 | 0.001 |
| museum_south_africa:museum_southeast_asia | squamates | 0.02736048 | 0.01060042 | 3.63286509 | 0.001 |
