## Supplemental Figures for "The role of recent speciation in present-day patterns of tetrapod phylogenetic relatedness"

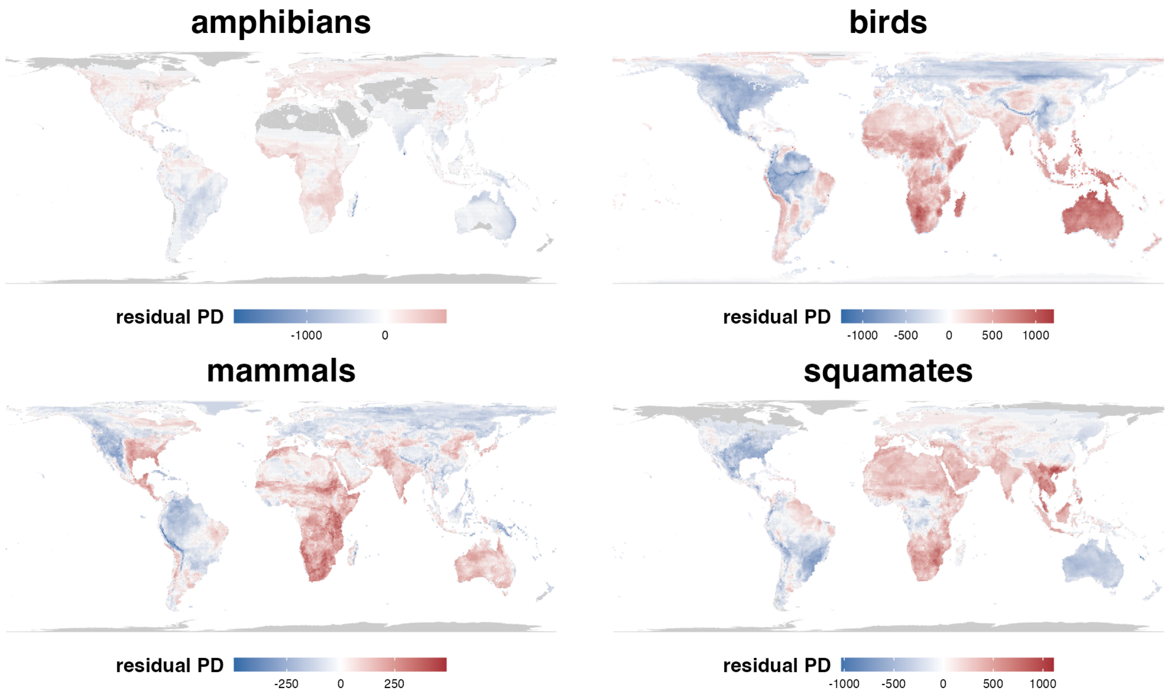


**Supplementary Figure 1**. Maps of residual phylogenetic diversity (PD) for all terrestrial vertebrate clades.

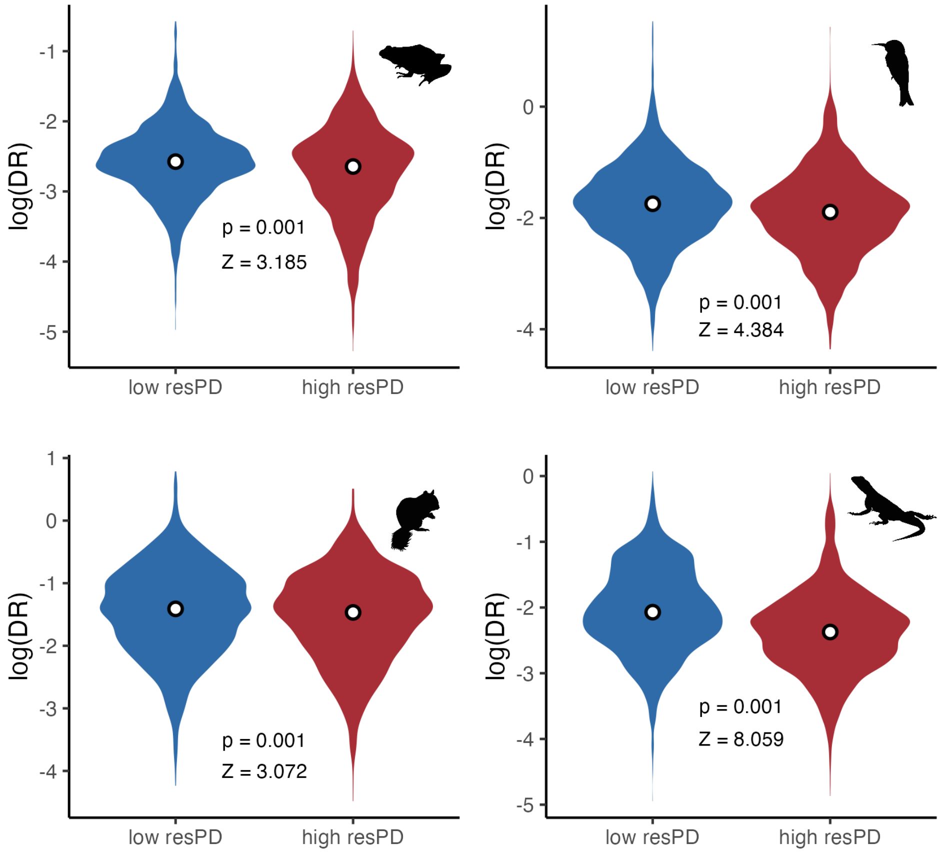


**Supplementary Figure 2**. Differences in recent speciation rates (DR rates) between areas of 10% lowest (in blue) and highest (in red) residual PD.

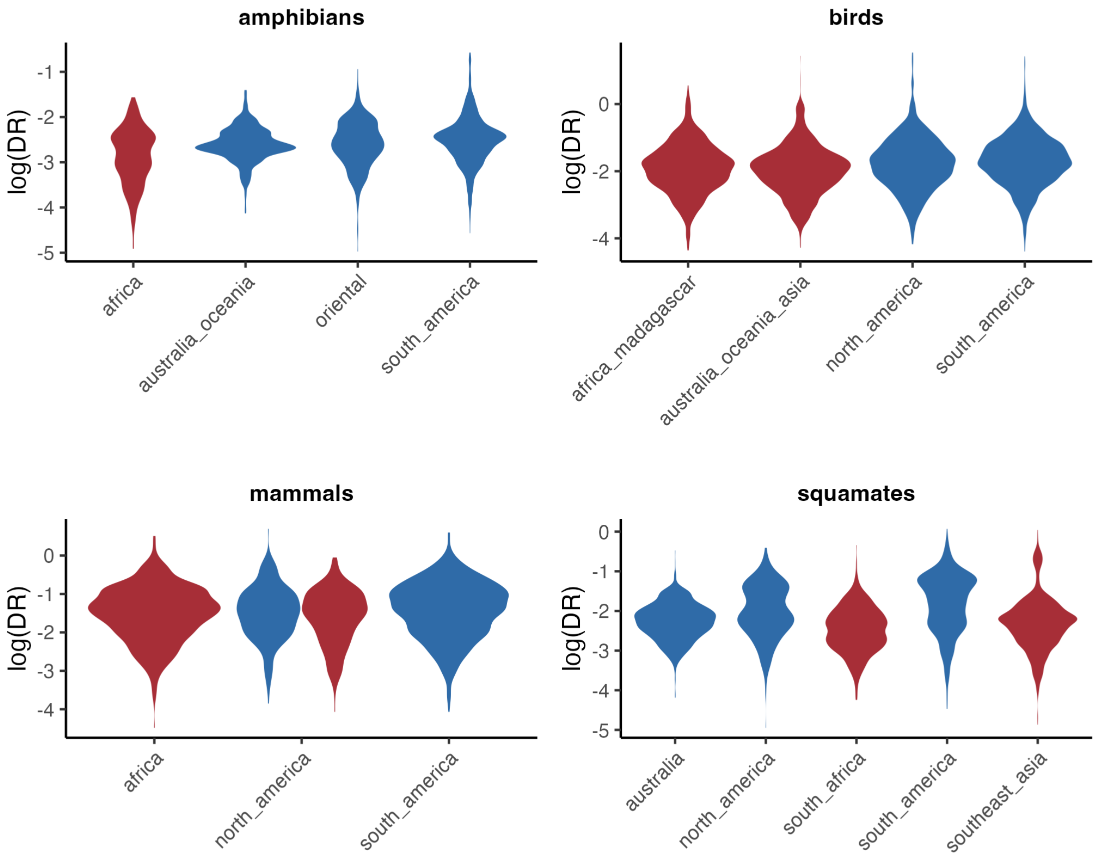


**Supplementary Figure 3**. Violin plots of recent speciation (DR) rates in the different focal areas identified as centers of high and low residual PD for all terrestrial vertebrate clades.

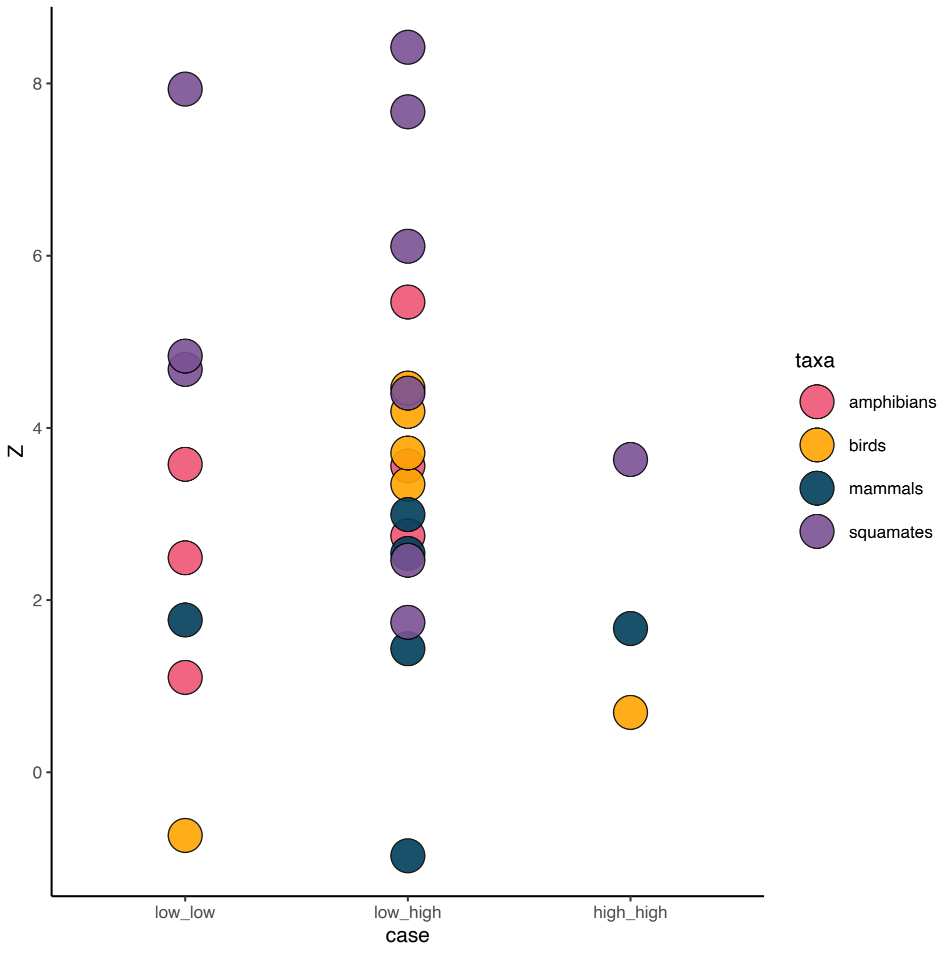


**Supplementary Figure 4**. Pairwise differences in DR rates (effect size Z) between specific regions of high and low residual PD, colored by taxa group. Left: differences between pairs of regions of low residual PD. Center: differences between regions of high residual PD and regions of low residual PD. Right: differences between pairs of regions of high residual PD.

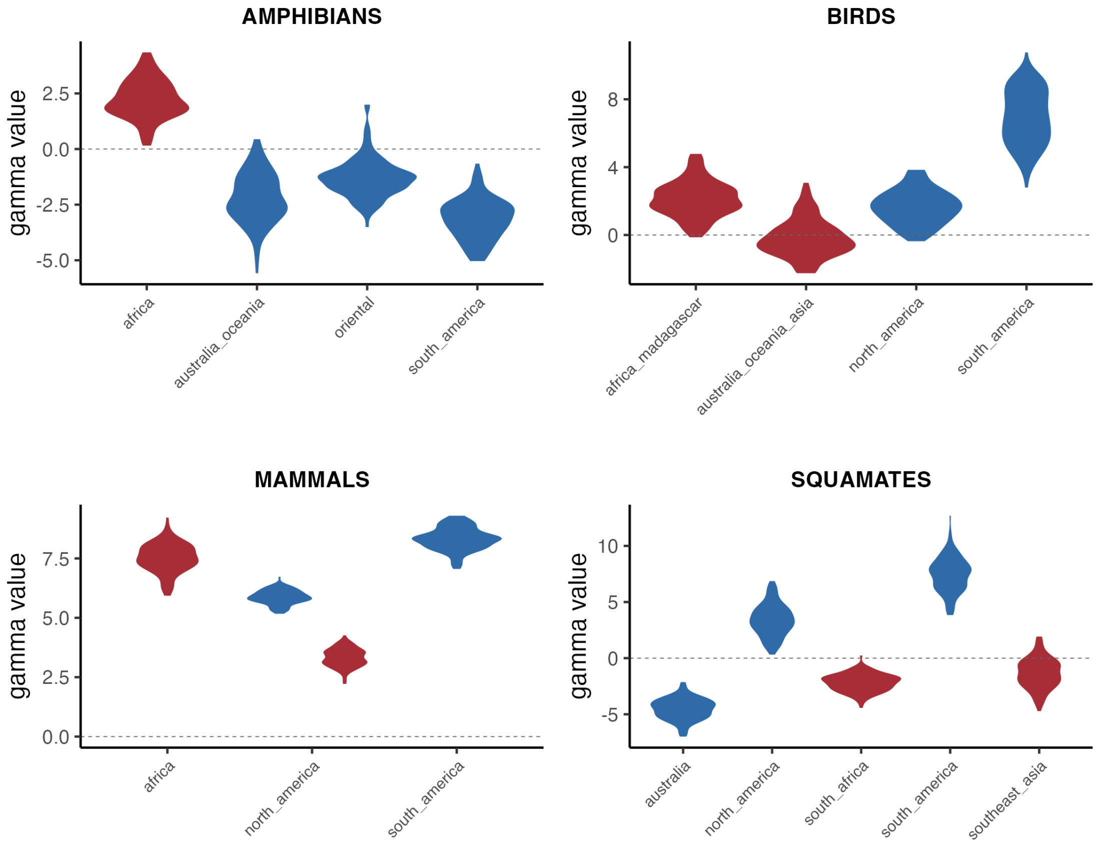


**Supplementary Figure 5**. Gamma-statistic in target regions of lowest and highest residual PD. Values above zero indicate nodes more concentrated toward the present compared to a constant birth-death model; values below zero indicate nodes concentrated toward the past.

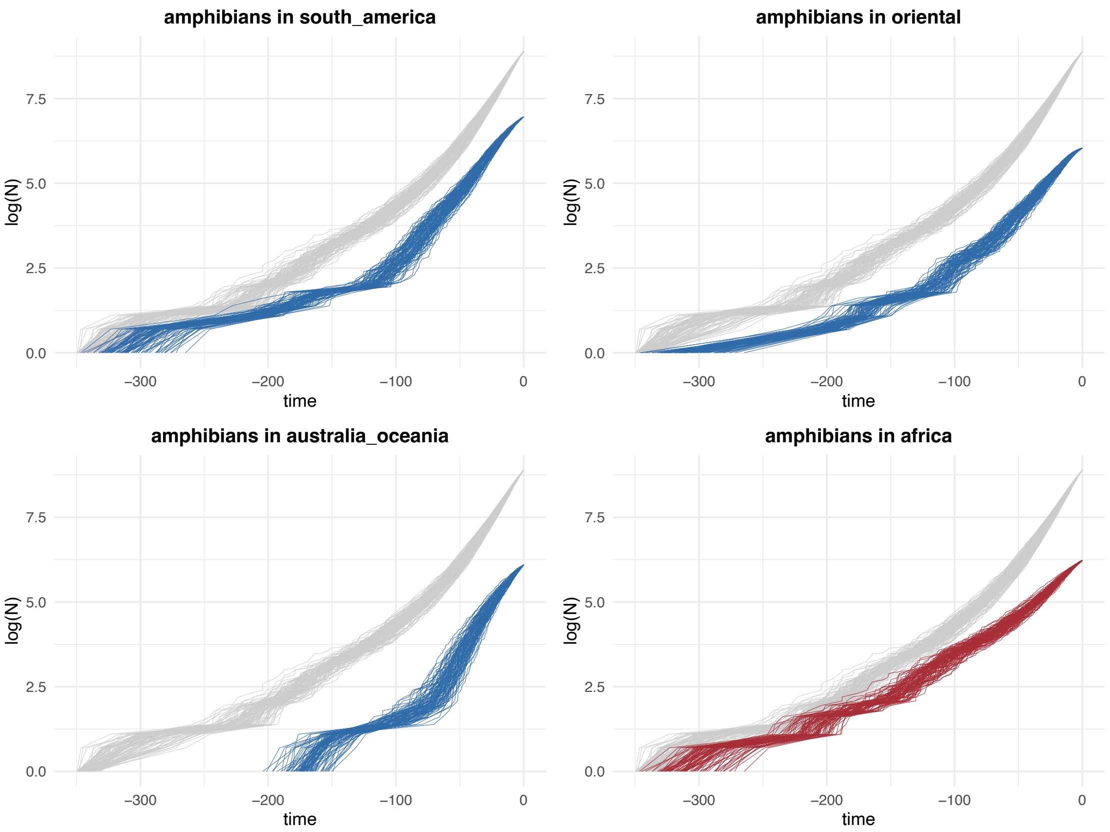


**Supplementary Figure 6**. Lineage-through-time (LTT) plots for amphibian lineages in the different focal regions of low (in blue) and high (in red) residual PD. In gray, LTT plot of the complete phylogeny for comparison.

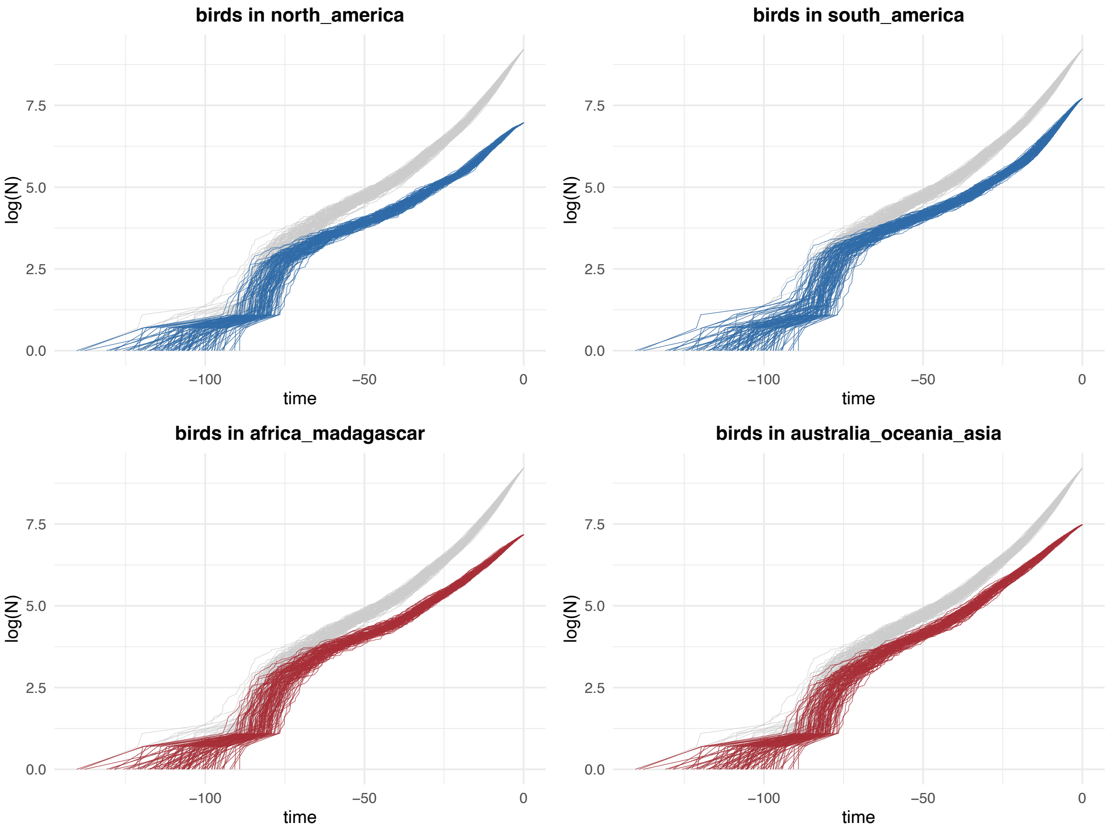


**Supplementary Figure 7**. Lineage-through-time (LTT) plots for bird lineages in the different focal regions of low (in blue) and high (in red) residual PD. In gray, LTT plot of the complete phylogeny for comparison.

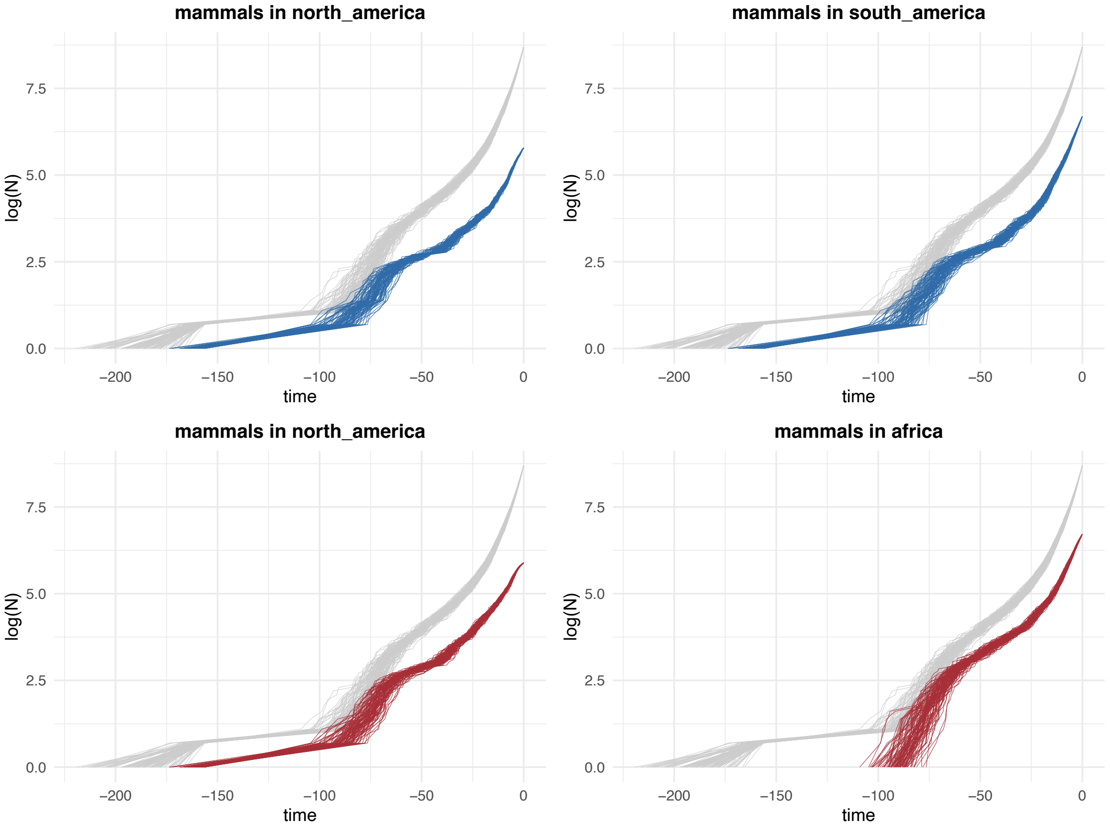


**Supplementary Figure 8**. Lineage-through-time (LTT) plots for mammal lineages in the different focal regions of low (in blue) and high (in red) residual PD. In gray, LTT plot of the complete phylogeny for comparison.

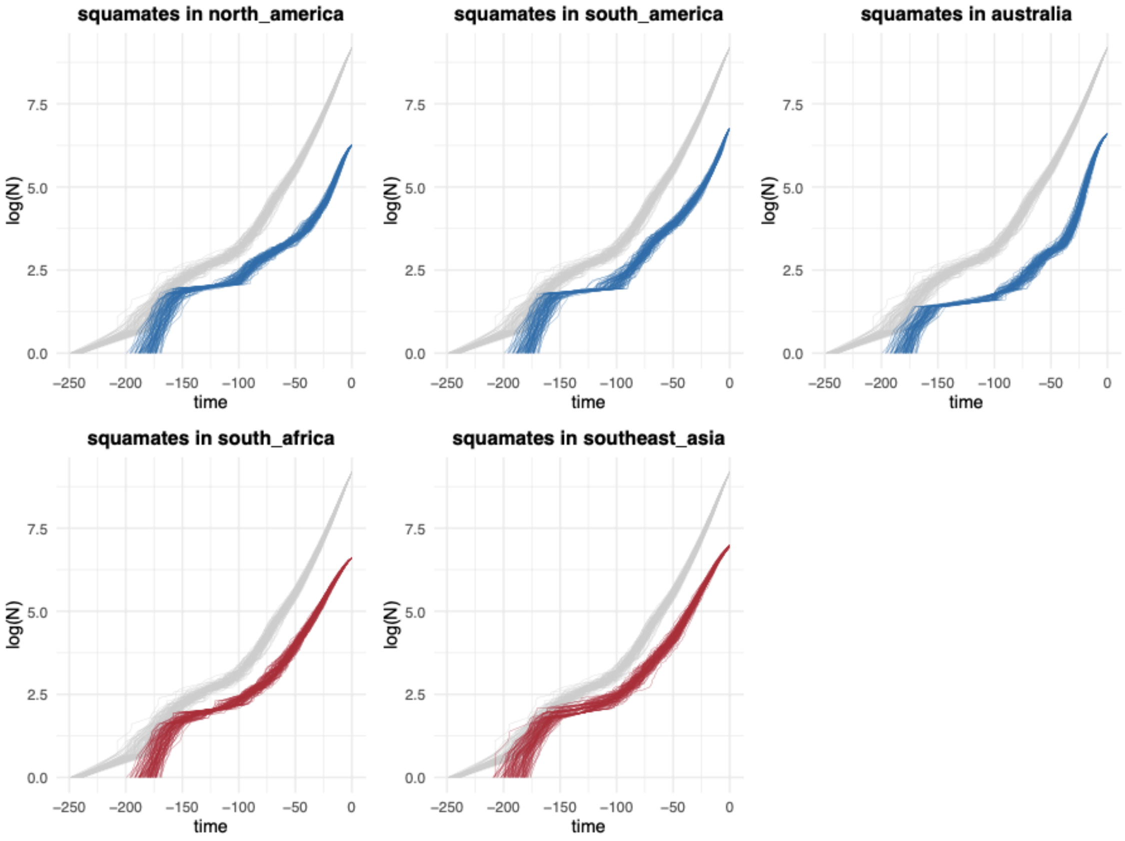


**Supplementary Figure 9**. Lineage-through-time (LTT) plots for squamate lineages in the different focal regions of low (in blue) and high (in red) residual PD. In gray, LTT plot of the complete phylogeny for comparison.

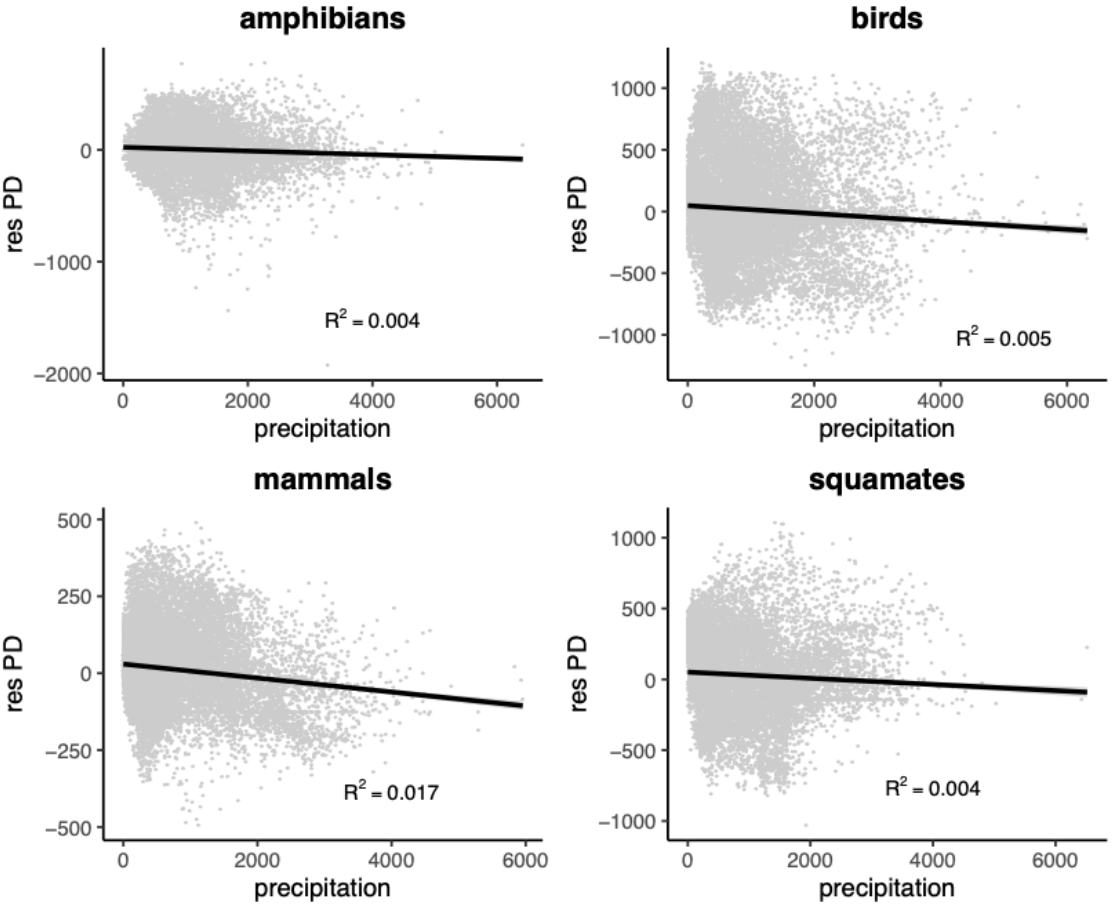


**Supplementary Figure 10**. Relationship of annual precipitation (in mm) with residual PD in all tetrapod clades.

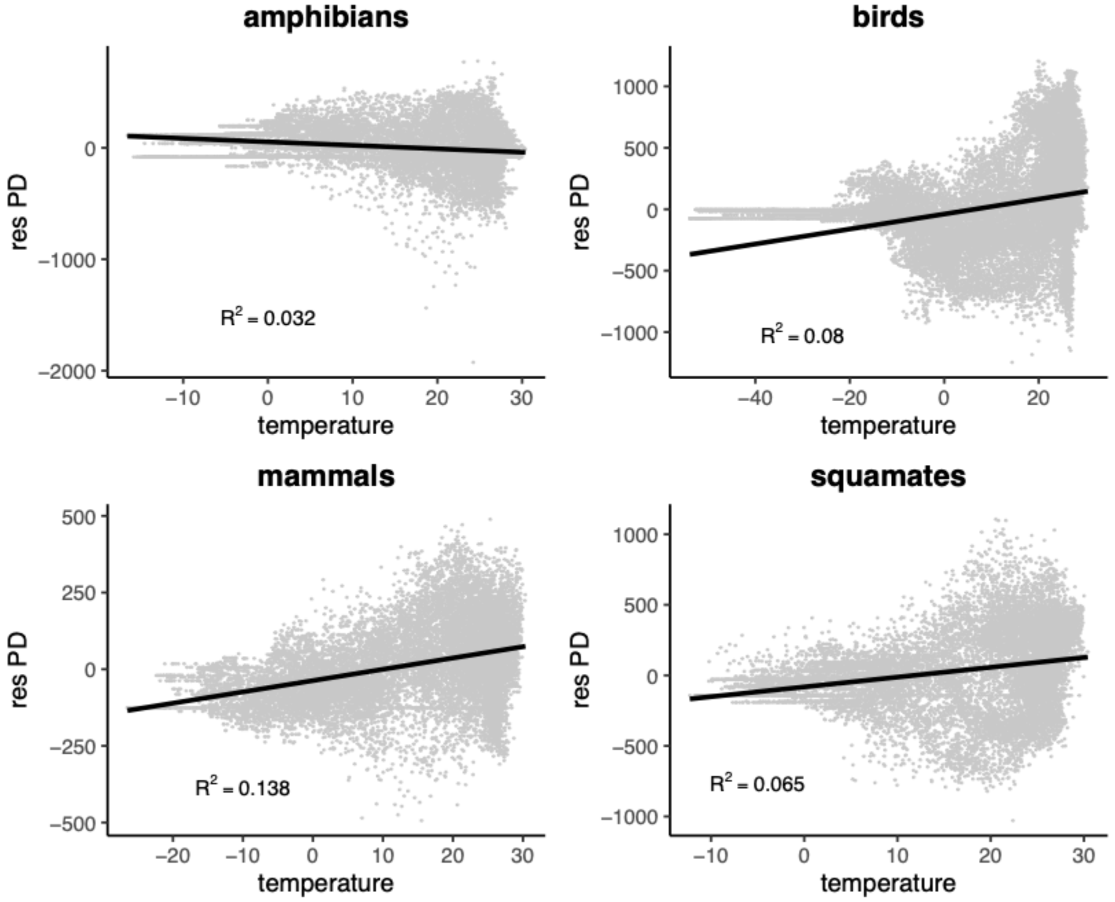


**Supplementary Figure 11**. Relationship of mean temperature (in ºC) with residual PD in all tetrapod clades.

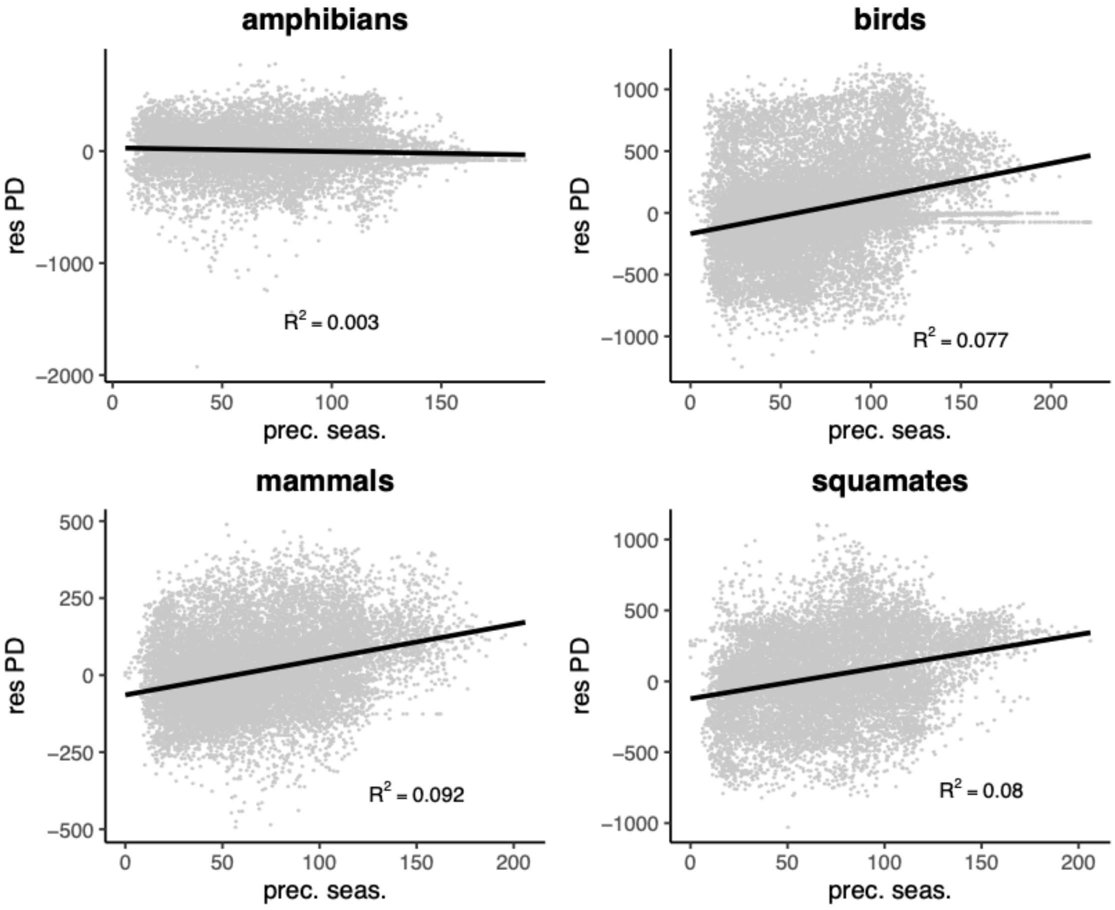


**Supplementary Figure 12**. Relationship of precipitation seasonality and residual PD in all tetrapod clades.

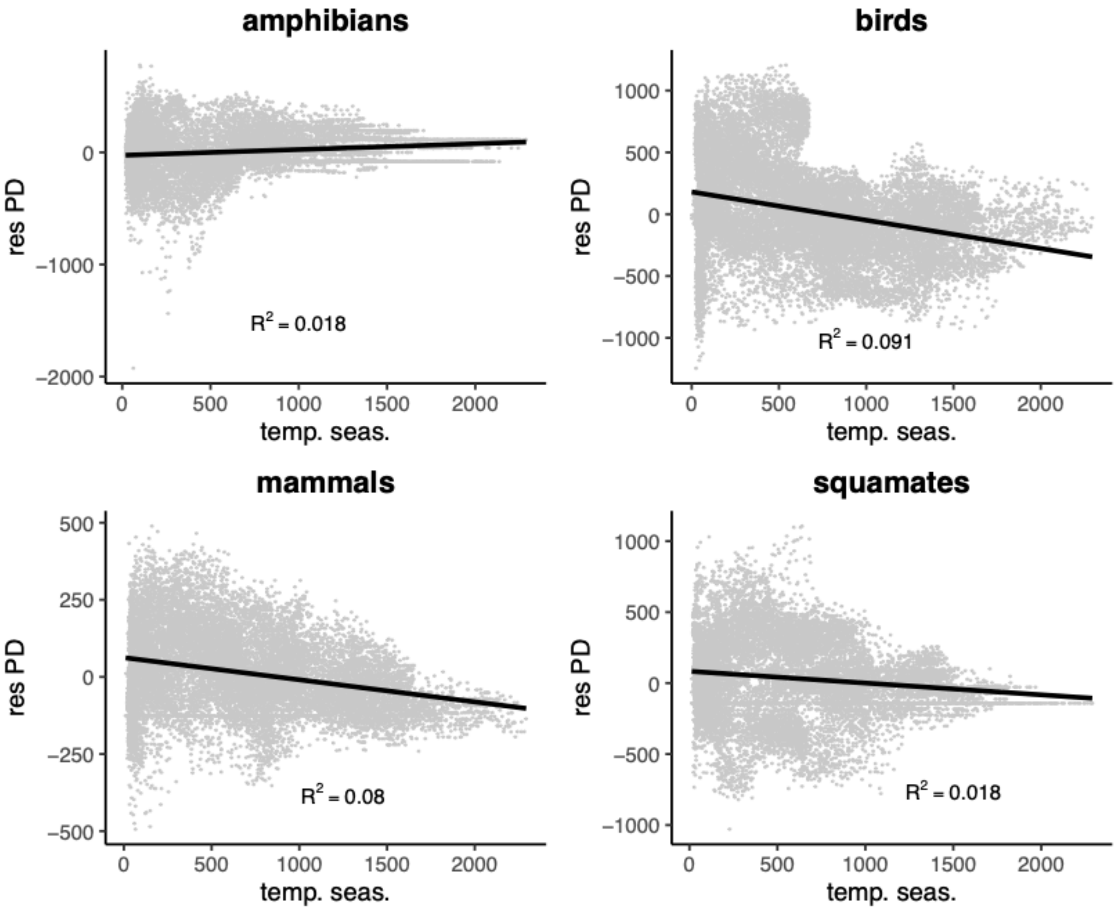


**Supplementary Figure 13**. Relationship of temperature seasonality and residual PD in all tetrapod clades.

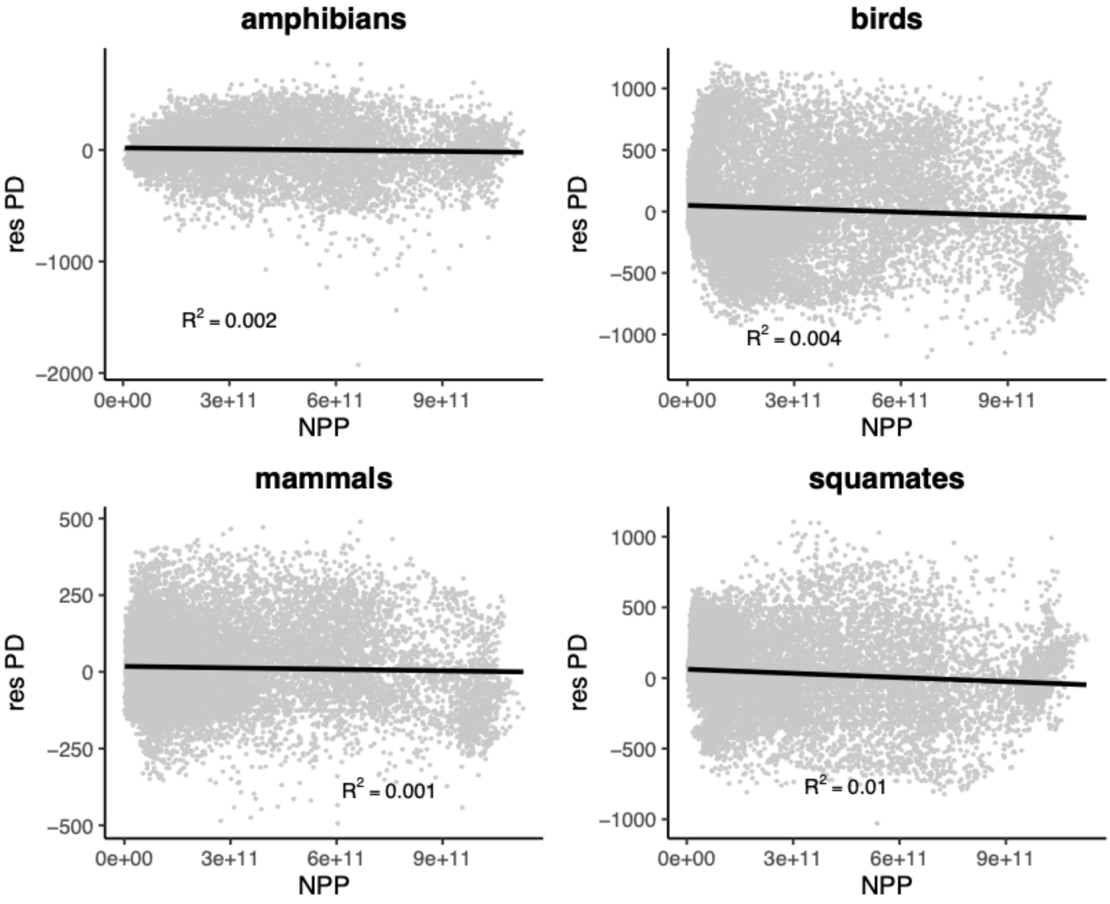


**Supplementary Figure 14**. Relationship of net primary productivity (NPP) and residual PD in all tetrapod clades.

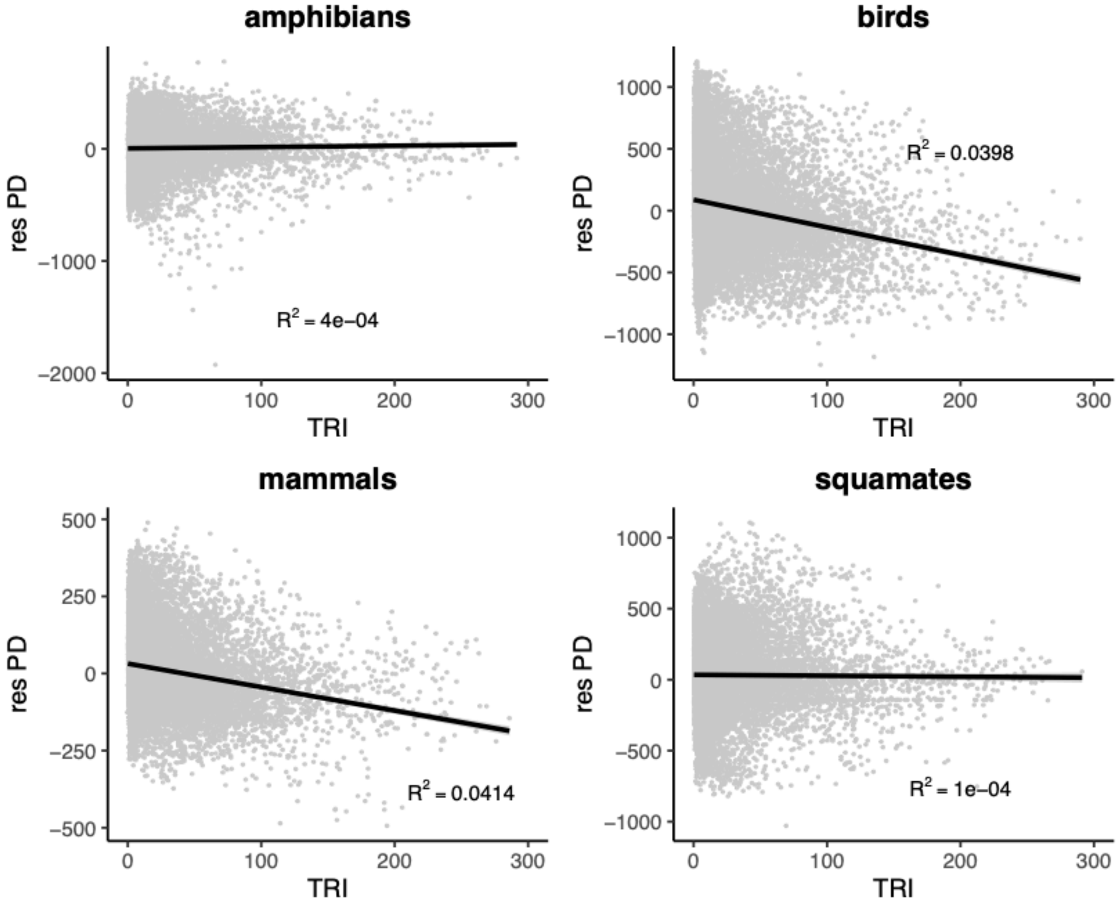


**Supplementary Figure 15**. Relationship of terrain roughness index (TRI; a measurement of topographic complexity) and residual PD in all tetrapod clades.

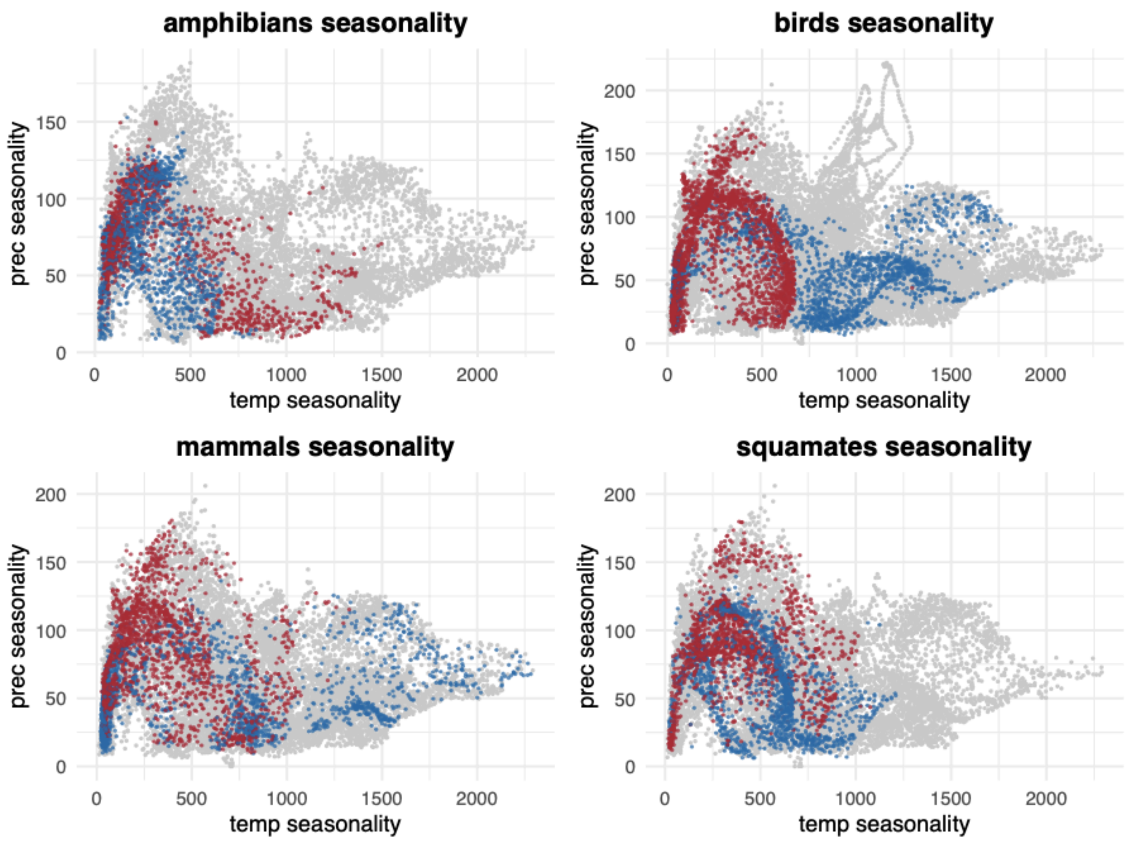


**Supplementary Figure 16**. Climatic space defined by precipitation seasonality and temperature seasonality for all tetrapod clades, showing the geographic grid cells with the 10% lowest (in blue) and highest (in red) residual PD.

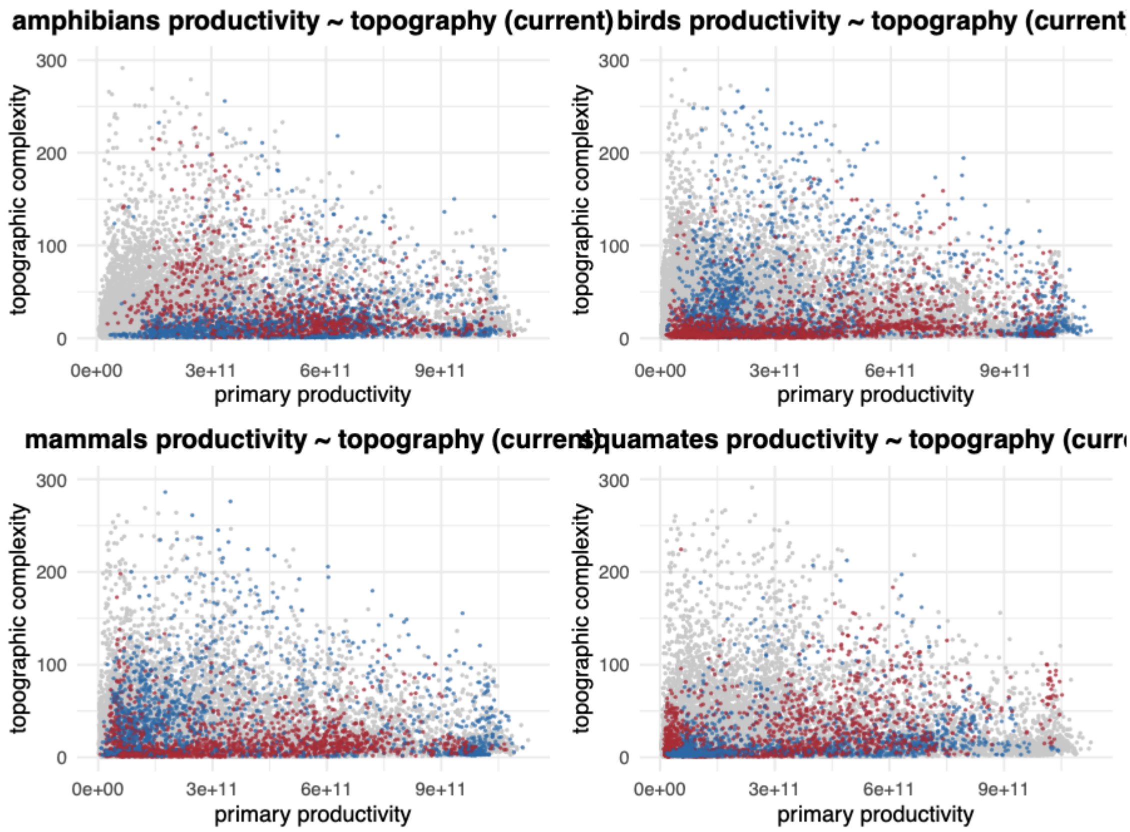


**Supplementary Figure 17**. Climatic space defined by net primary productivity (NPP) and terrain roughness index (TRI) for all tetrapod clades, showing the geographic grid cells with the 10% lowest (in blue) and highest (in red) residual PD.

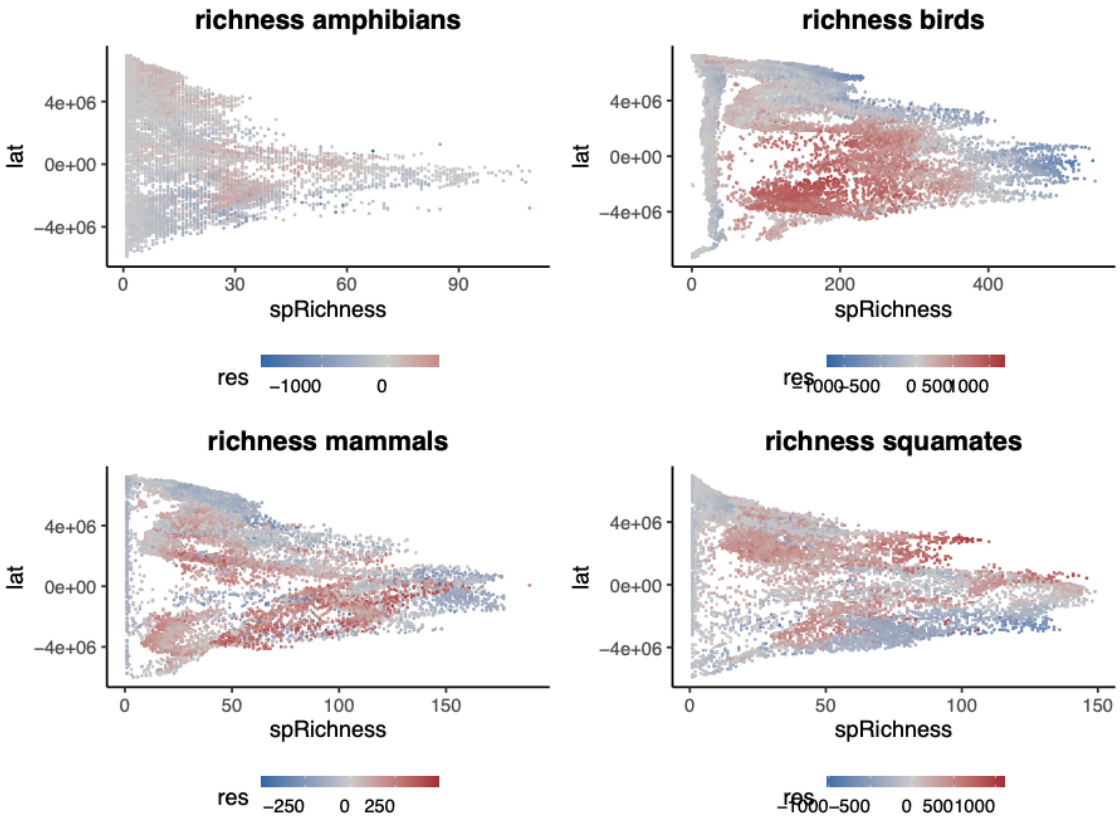


**Supplementary Figure 18**. Latitudinal pattern of species richness (in the X-axis) and residual PD (in the color gradient) for all tetrapod clades.
