## Appendix I for "The role of recent speciation in present-day patterns of tetrapod phylogenetic relatedness"

### Multiple regression models: temperature and precipitation

#### Load packages

```r
# packages ----
packages <- c('RRPP')
easypackages::libraries(packages)
```

#### Load data

```r
# hexagonal grid vertebrates ----
hexgrid_list <- readRDS('../../objects/hexgrid_list_geo_v3.rds')
taxa <- names(hexgrid_list)
```

```r
# see the data (e.g., for amphibians)
head(hexgrid_list[['amphibians']])
```

```
##      spRichness         dr     logdr       pd   id      group resloess_pd_rich
## 676           1 0.07100814 -2.644961 311.5848  676 amphibians         -82.7005
## 752           1 0.07100814 -2.644961 311.5848  752 amphibians         -82.7005
## 827           1 0.07100814 -2.644961 311.5848  827 amphibians         -82.7005
## 903           1 0.07100814 -2.644961 311.5848  903 amphibians         -82.7005
## 978           1 0.07100814 -2.644961 311.5848  978 amphibians         -82.7005
## 1054          1 0.07100814 -2.644961 311.5848 1054 amphibians         -82.7005
##      type  geo        temp      prec  tempseas precseas          npp
## 676  none <NA>  1.33583746 1057.6309  681.5326 31.87319  36663643788
## 752  none <NA> -3.34670127  362.4686 1131.8237 55.57872  92041091651
## 827  none <NA> -0.17327846 1352.6915  764.5944 29.75463  52426766080
## 903  none <NA> -3.32864311  405.1696 1153.3876 50.80284 107454674893
## 978  none <NA>  0.04908044 1558.8645  799.4727 28.22991  88986482495
## 1054 none <NA> -5.52950498  563.3844 1022.5608 49.02254  89259016281
##      tri_current tri_holo  tri_lgm      long     lat
## 676     98.38327 81.27259 69.28321 -14130128 6396996
## 752     38.39680 38.39680 38.39680 -14080128 6483598
## 827    101.85556 91.80629 86.84954 -14030128 6396996
## 903     45.87827 45.87827 45.87827 -13980128 6483598
## 978     99.05036 95.77317 94.36873 -13930128 6396996
## 1054    94.60183 94.60183 94.60183 -13880128 6483598
```

### Analysis

```
# Residual PD vs temp*prec ----
# LM res~temp*prec
fit.res_temprec <- vector('list', length(taxa))
rsq_res_temprec <- vector('numeric', length(taxa))
names(rsq_res_temprec) <- names(fit.res_temprec) <- taxa
for (t in taxa){
  fit.res_temprec[[t]] <- lm.rrpp(resloess_pd_rich ~ temp*prec, SS.type = "III",
                                  data = hexgrid_list[[t]])
  fit.sum <- summary(fit.res_temprec[[t]])
  fit.sum$table$Rsq
  rsq_res_temprec[t] <- fit.sum$table$Rsq

}

# see Rsq values
lapply(fit.res_temprec, summary)
```

```
## $amphibians
##
## Linear Model fit with lm.rrpp
##
## Number of observations: 12615
## Number of dependent variables: 1
## Data space dimensions: 1
## Sums of Squares and Cross-products: Type III
## Number of permutations: 1000
##
## Full Model Analysis of Variance
##
##             Df Residual Df       SS Residual SS        Rsq        F Z (from F)
## temp * prec  3       12611 20929686   483160708 0.04151971 182.0956   15.03156
##             Pr(>F)
## temp * prec  0.001
##
##
## Redundancy Analysis (PCA on fitted values and residuals)
##
##              Trace Proportion Rank
## Fitted     1659.24  0.0415196    1
## Residuals 38303.53  0.9584803    1
## Total     39962.77  1.0000000    1
##
## Eigenvalues
##
##                PC1
## Fitted     1659.24
## Residuals 38303.53
## Total     39962.77
##
##
## $birds
```

```
##
## Linear Model fit with lm.rrpp
##
## Number of observations: 19750
## Number of dependent variables: 1
## Data space dimensions: 1
## Sums of Squares and Cross-products: Type III
## Number of permutations: 1000
##
## Full Model Analysis of Variance
##
##              Df Residual Df        SS Residual SS       Rsq        F Z (from F)
## temp * prec  3       19746 437532337  2275272230 0.1612841 1265.711   20.05509
##              Pr(>F)
## temp * prec  0.001
##
##
## Redundancy Analysis (PCA on fitted values and residuals)
##
##                Trace Proportion Rank
## Fitted     22154.66  0.1612842    1
## Residuals 115209.49  0.8387159    1
## Total     137364.15  1.0000000    1
##
## Eigenvalues
##
##                  PC1
## Fitted     22154.66
## Residuals 115209.49
## Total     137364.15
##
##
## $mammals
##
## Linear Model fit with lm.rrpp
##
## Number of observations: 15747
## Number of dependent variables: 1
## Data space dimensions: 1
## Sums of Squares and Cross-products: Type III
## Number of permutations: 1000
##
## Full Model Analysis of Variance
##
##              Df Residual Df       SS Residual SS       Rsq       F Z (from F)
## temp * prec  3       15743 59124775   203630068 0.2250188 1523.68   20.85565
##              Pr(>F)
## temp * prec  0.001
##
##
## Redundancy Analysis (PCA on fitted values and residuals)
##
##                Trace Proportion Rank
## Fitted     3754.908  0.2250188    1
```

```
## Residuals 12932.178  0.7749812    1
## Total     16687.085  1.0000000    1
##
## Eigenvalues
##
##                 PC1
## Fitted     3754.908
## Residuals 12932.178
## Total     16687.085
##
##
## $squamates
##
## Linear Model fit with lm.rrpp
##
## Number of observations: 13961
## Number of dependent variables: 1
## Data space dimensions: 1
## Sums of Squares and Cross-products: Type III
## Number of permutations: 1000
##
## Full Model Analysis of Variance
##
##             Df Residual Df        SS Residual SS        Rsq        F Z (from F)
## temp * prec  3       13957 104383285  1004309641 0.09414986 483.5419   19.56422
##             Pr(>F)
## temp * prec  0.001
##
##
## Redundancy Analysis (PCA on fitted values and residuals)
##
##              Trace Proportion Rank
## Fitted     7477.31  0.0941498    1
## Residuals 71941.95  0.9058501    1
## Total     79419.26  0.9999999    1
##
## Eigenvalues
##
##                PC1
## Fitted     7477.31
## Residuals 71941.95
## Total     79419.26
```

```
# see significance of different terms
lapply(fit.res_temprec, anova)
```

```
## $amphibians
##
## Analysis of Variance, using Residual Randomization
## Permutation procedure: Randomization of null model residuals
## Number of permutations: 1000
## Estimation method: Ordinary Least Squares
## Sums of Squares and Cross-products: Type III
## Effect sizes (Z) based on F distributions
```

```
##
##              Df        SS      MS     Rsq       F      Z Pr(>F)
## temp          1    513296  513296 0.00102  13.398 2.8454  0.001 **
## prec          1   4688390 4688390 0.00930 122.372 6.1058  0.001 **
## temp:prec     1   4706327 4706327 0.00934 122.840 6.3158  0.001 **
## Residuals 12611 483160708   38313 0.95848
## Total     12614 504090395
## ---
## Signif. codes:  0 '***' 0.001 '**' 0.01 '*' 0.05 '.' 0.1 ' ' 1
##
## Call: lm.rrpp(f1 = resloess_pd_rich ~ temp * prec, SS.type = "III",
##     data = hexgrid_list[[t]])
##
## $birds
##
## Analysis of Variance, using Residual Randomization
## Permutation procedure: Randomization of null model residuals
## Number of permutations: 1000
## Estimation method: Ordinary Least Squares
## Sums of Squares and Cross-products: Type III
## Effect sizes (Z) based on F distributions
##
##              Df         SS        MS     Rsq       F      Z Pr(>F)
## temp          1   94108675  94108675 0.03469  816.72 10.693  0.001 **
## prec          1  186696465 186696465 0.06882 1620.25 12.890  0.001 **
## temp:prec     1  103323311 103323311 0.03809  896.69 10.536  0.001 **
## Residuals 19746 2275272230    115227 0.83872
## Total     19749 2712804567
## ---
## Signif. codes:  0 '***' 0.001 '**' 0.01 '*' 0.05 '.' 0.1 ' ' 1
##
## Call: lm.rrpp(f1 = resloess_pd_rich ~ temp * prec, SS.type = "III",
##     data = hexgrid_list[[t]])
##
## $mammals
##
## Analysis of Variance, using Residual Randomization
## Permutation procedure: Randomization of null model residuals
## Number of permutations: 1000
## Estimation method: Ordinary Least Squares
## Sums of Squares and Cross-products: Type III
## Effect sizes (Z) based on F distributions
##
##              Df        SS       MS     Rsq         F       Z Pr(>F)
## temp          1  34827331 34827331 0.13255 2692.5624 13.0002  0.001 **
## prec          1     21455    21455 0.00008    1.6587  0.8590  0.218
## temp:prec     1   1905952  1905952 0.00725  147.3525  7.1825  0.001 **
## Residuals 15743 203630068    12935 0.77498
## Total     15746 262754843
## ---
## Signif. codes:  0 '***' 0.001 '**' 0.01 '*' 0.05 '.' 0.1 ' ' 1
##
## Call: lm.rrpp(f1 = resloess_pd_rich ~ temp * prec, SS.type = "III",
##     data = hexgrid_list[[t]])
```

```
##
## $squamates
##
## Analysis of Variance, using Residual Randomization
## Permutation procedure: Randomization of null model residuals
## Number of permutations: 1000
## Estimation method: Ordinary Least Squares
## Sums of Squares and Cross-products: Type III
## Effect sizes (Z) based on F distributions
##
##               Df         SS       MS     Rsq      F      Z Pr(>F)
## temp           1   16192989 16192989 0.01461 225.04 7.5257  0.001 **
## prec           1   14179876 14179876 0.01279 197.06 7.1941  0.001 **
## temp:prec      1    6895668  6895668 0.00622  95.83 5.8765  0.001 **
## Residuals 13957 1004309641    71957 0.90585
## Total     13960 1108692926
## ---
## Signif. codes:  0 '***' 0.001 '**' 0.01 '*' 0.05 '.' 0.1 ' ' 1
##
## Call: lm.rrpp(f1 = resloess_pd_rich ~ temp * prec, SS.type = "III",
##     data = hexgrid_list[[t]])
```
